# Gonadal Sex and Sex-Chromosome Complement Interact to Affect Ethanol Consumption in Adolescent Four Core Genotypes Mice

**DOI:** 10.1101/2022.10.25.513748

**Authors:** Shawn M. Aarde, Jared R. Bagley, J. David Jentsch

**Author notes:** Aarde (corresponding author).

## Abstract

**Background:** Sex differences in ethanol consumption have been reported in both humans and laboratory rodents, but the independent/dependent contributions of genetic and hormonal sex◻biasing mechanisms to these phenotypes have not yet been fully explored.

**Methods:** To examine the contributions of sex-chromosome complement (SCC) and gonadal sex (GS) to ethanol consumption, we studied adolescent (28-32 days old) four core genotypes (FCG) mice (C57BL/6J background; FCG model allows for independent assortment of GS and SCC) using a modified drinking-in-the-dark (DID) procedure. Mice were offered concurrent access to 20%, 10% and 0% ethanol (in water) in four daily 2-hour sessions. Consumption at the level of individual bouts was recorded.

**Results:** Although all four genotype groups preferred the 20% ethanol over 10% and 0%, and showed similar consumption of the 10% and 0% solutions, the group rankings for consumption of the 20% ethanol solution were XX+testes > XY+testes > XY+ovaries > XX+ovaries. Thus, an interaction was observed between SCC and GS for which the simple effect of SCC was greatest in mice with ovaries (XY > XX) and the simple effect of GS was greatest in XX mice (testes > ovaries). Moreover, these effects varied in magnitude across and within drinking sessions. The behavioral microstructure of ethanol consumption (i.e., parameterization of within-session discriminable drinking bouts) support the validity of our 3-bottle modification of the DID procedure as a model of binge-like consumption as: (1) the consumption rate of the 20% ethanol solution was ~80 g EtOH/kg/h *within a bout* (~12 s/bout, ~3 bouts/session), (2) most of this ethanol consumption was completed in a single bout and (3) within-session ethanol consumption was greater earlier than later, indicating “front loading.”

**Conclusions:** These results indicate that SCC and GS interact on ethanol consumption in adolescent FCG mice on a C57BL/6J background to affect binge-like consumption from the very initiation of access and that these effects are dynamic as they varied both across and within sessions.

**Highlights:**

- Gonadal sex and sex-chromosome complement *interact* on ethanol consumption in adolescent four core genotypes mice
- In adolescent four core genotypes mice, mice with testes drink more ethanol than mice with ovaries, particularly in the presence of an XX karyotype
- In adolescent four core genotypes mice, XY mice drink more ethanol than XX mice, but only in mice with ovaries
- The effects of sex-biasing biological factors on the patterns of ethanol consumption by adolescent four core genotypes mice that we observed in our 3-bottle Drinking-in-the-Dark procedure showed face validity with some of the sex/gender differences observed in human adolescents

## BACKGROUND

Biological sex has been associated with differences in patterns of adolescent alcohol (ethanol) consumption (e.g., binge frequency, age of first experience and rate of progression, motivations to drink, and reactivity to positive/negative experiences) [1–10], as well as to alcohol use disorder (AUD) [1–3,5,10,11]. These observations implicate effects of, and interactions between, gonadal hormones [12–18], gene-sex interactions [19–24] and/or gendered socio-cultural factors [7,25,26] on responsivity to ethanol.

Of particular concern is adolescent binge drinking, which is defined as the consumption of enough ethanol within 2 hours to achieve a blood ethanol concentration (BEC) ≥ 80 mg/dL [27] as it predicts future alcohol dependence and alcohol use disorder (AUD) [1,28–32]. It has been estimated that ~77% of the costs associated with alcohol use in the U.S. are attributable to binge drinking [33]. Sex differences in alcohol drinking increase from 8^th^ to 12^th^ grade by which stage bingeing is substantially greater in males (16%) than females (12%) [6,34]. Although this sex difference continues into adulthood, its magnitude has been decreasing over time largely due to increasing alcohol use by females [35–37].

Sex differences in phenotypes like binge drinking could be attributable to three key sex-biasing factors: organizational and activational mechanisms (mediated by gonadal type and associated steroid secretions), as well as to sex-chromosome complement (SCC) mechanisms (mediated by differences in X and Y genes and their expression dosage) [38]. People with an XY sex◻chromosome complement typically have a functional SRY (sex determining region of the Y-chromosome) gene that guides gonad differentiation toward the development of testes, or *sans* SRY toward the development of ovaries, which subsequently determines the type of gonadal hormones secreted [39,40]. These coupled mechanisms make it difficult to fully dissect the causal effects of, and interactions between, these three sex-biasing factors.

However, animal models are available that enable decoupling the *Sry* gene from the Y-chromosome. For example, in the four core genotypes (FCG) mouse, the Y chromosome has been deleted for *Sry* which is instead expressed from an autosomal transgene [41–44]), thereby making gonadal sex (GS) independent from SCC. Research using this model has shown direct genetic effect of SCC on ethanol consumption [24] and on the formation of habitual operant responding for ethanol [19]. Moreover. the importance of both the direct effects of SCC and its interactions with GS on basic reward/reinforcement processes, as well as cognitive functions that are centrally implicated in AUD and other addictions, has been underscored by both human studies on sex-chromosome aneuploidies [45,46] and in animal models like the FCG mouse [47–52,52,53] and those that model sex-chromosome aneuploidies [54–61].

Thus, discovering and validating useful models of sex-chromosome effects on alcohol consumption and AUD would greatly benefit from testing a preclinical animal model like the FCG mouse using a procedural model of ethanol-bingeing behavior. One such model that has demonstrated construct and predictive validity in the background strain of the FCG mouse – C57BL/6J – is the drinking-in-the-dark (DID) model established by Rhodes et al [62–69]. The key feature of this procedure is the circadian placement and duration of ethanol access; mice are provided limited (2-4 h) access to an ethanol solution (typically, 20%) in their home cage starting 3 hours after the onset of the animals’ dark-phase. Binge-like ethanol intake was confirmed by BECs for which mice regularly exceed values of 100 mg/dL. Moreover, studies of DID that include behavioral microstructural intake data (i.e., parameterized high temporal resolution behavioral recordings) have similarly shown that mice concentrate ethanol consumption into short discriminable binge-like bouts [66,70,71] and that sex differences can be detected in both total intake and within-session changes in intake (e.g., “front loading”) [72].

Importantly, although sex differences observed in DID-based procedures typically show greater ethanol consumption in female mice than male mice [73–79], these sex differences are sometimes not detected [80–84], varied from one experiment to the next within a report [66,67,72,85] and/or show greater intake in males than females [71]. This suggests that some procedural factors that vary across research studies moderate sex effects on ethanol drinking behaviors.

Additionally, using a non-DID drinking procedure, Sneddon et al studied FCG mice [24] that were provided access to increasing concentrations of ethanol (5% → 10% → 15% → 20%; clusters of 5 consecutive sessions/concentration at 24 h/session) and water. Group differences in ethanol consumption were concentration dependent, as XX mice consumed more ethanol than XY mice for only the 15% solution (similarly for preference; XX > XY only at 15% ethanol), and mice with ovaries consumed more ethanol than mice with testes for only the 20% solution (no GS differences in preference were detected). Thus, one way to maximize the likelihood of detecting the effects of SCC, GS and their interaction in the DID procedure may also be to include access to a range of concentrations.

Based on the aforementioned literature, we sought to examine ethanol intake and preference in adolescent, gonad-intact FCG mice. Thus, we used a DID procedure that offered mice access to water vs. two concentrations of ethanol (10 and 20%). We hypothesized that we would find that mice with ovaries would drink more ethanol than mice with testes and would do so by engaging in smaller, but more frequent, consumption bouts [70,71]. Additionally, based on the only currently available FCG mouse studies of ethanol drinking (one using inbred mice on the C57BL/6J background [24] and the other on outbred mice of the MF1 background [19]), we additionally predicted greater ethanol consumption by XX mice than XY mice [24], as well as potentially an attenuated interaction whereby the SCC effect (XX > XY) would be greater in mice with testes than in those with ovaries [24] and the GS effect (mice with ovaries (**O**) > mice with testes (**T**)) would be greater in XX mice [19].

## METHODS

### Subjects

#### FCG mouse model

Subjects were early adolescent mice (28-29 days old at DID start) from the FCG model [86]; FCG mice were originally maintained on an MF1 background before full backcrossing to the C57BL/6J genetic background. In the FCG model, the Y chromosome is deleted for *Sry* (called the Y^−^ chromosome), and *Sry* is introduced onto chromosome 3 as a transgene (*Sry*+) [44]. XY^−^(*Sry+*) sires mated with XX(*Sry*-) dams produced progeny of four genotypes: XY^−^(*Sry*+) with testes, XX(*Sry*+) with testes, XY^−^(*Sry*−) with ovaries and XX(*Sry*−) with ovaries. For simplicity, with reference to gonadal sex (**GS**) in FCG mice, we describe *Sry*-mice as mice with ovaries (**O**) and *Sry*+ mice as mice with testes (**T**). Thus, we refer to the four core genotypes as **XX+O** (XX+ovaries; N=39), **XX+T** (XX+testes; N=39), **XY+O** (XY+ovaries; N=38) and **XY+T** (XY+testes; N=35). Genotypes were determined via standardized probes for both Y^−^ and *Sry* via genotyping services provided by Transnetyx (Transnetyx, Inc., NY, USA) and confirmed by gonadal phenotype (Burgoyne and Arnold, 2016). Procedures and protocols were approved by the Institutional Animal Care and Use Committee (IACUC) at Binghamton University and were carried out in a manner consistent with the “Guide for the Care and Use of Laboratory Animals, Eighth Edition” [87].

#### Husbandry

Mice were group housed (2-4 mice/cage of the same GS) in standard open-top home cages with woodchip bedding (to minimized phytoestrogens/pseudoestrogens found in corncob bedding [88]), *ad libitum* chow (PicoLab^®^ Laboratory Rodent Diet 5LOD) [89], and environmental enrichment that included a nesting pack (~2”x2” paper pillow stuffed with strips of paper), a small wood block and a red plastic PVC tube. Subjects were kept under a 12:12 light:dark cycle in a temperature (~72 °F) and humidity (~46%) controlled vivarium.

### Drinking-in-the-Dark Procedure

#### Overview, Equipment

Liquid (LIQ) consumption was monitored via a 3-bottle apparatus attached to a modified standard home cage as described previously [90]. Bottles sit on a load cell that measures small changes in bottle weight, and contact by the animal, at a sampling rate of ~2.2 samples/second. Thus, load cells serve as contact sensors and mass scales that permit high resolution discrimination of both bout frequency/duration and amount consumed per bout. Bottles were constructed from modified 50 mL conical tubes fitted with a silicone stopper and curved sipper tube within which were two steel ball bearings to minimize non-contact liquid loss. As the temporally discrete recording of consumption includes a taring and re-taring of bottle weight, the contamination of non-contact liquid loss (e.g., “leak”) is minimized [90]. Test cages were placed within sound and light attenuating cubicles equipped with a fan for ventilation and white noise. Each cage contained a fresh layer of woodchip bedding and were topped with a standard wire cage lid.

#### Habituation Phase

On the day prior to the first DID session, mice were habituated to handling (weighing, tail marking), transport to the testing room, the testing room environment (all electronic equipment on), and ethanol odor/taste by dropping ~1 ml of 20% ethanol on their home cage nesting pack.

#### Drinking in the Dark (DID) Testing Phase

The day after habituation, mice were tested for liquid consumption over four sequential 2-h DID sessions that began 2 h after the start of the dark phase of their light cycle. Three bottles of different ethanol concentrations (20%, 10% and 0%; in tap water) were provided, as was a ~4 gram food pellet. Two to 3 hours before each session, mice were weighed, tail marked and transported to the testing room. Mice were weighed again immediately after each session and returned to the vivarium within 30 min.

#### Blood Ethanol Concentrations (BEC)

To estimate the approximate BECs of mice immediately after DID sessions, serum from trunk blood was collected from a subset of mice within 5 min of the end of the last DID session and stored at −80° C until being analyzed with the Analox system (Analox Instruments Ltd, UK; AM1 Alcohol Analyser using a 100 mg/dL calibration standard and available quality control serum [GMRD-110D4]). This group included mice from all 4 genotypes (XX+**O** N=11, XY+**O** N=6, XX+**T** N=7, XY+**T** N=9).

### Measures, Design and Data Analyses

Behavior was recorded as a sequence of discriminable bouts of liquid consumption as described previously [90]. To filter out spout contacts that were not associated with drinking, bouts were delimited by pauses of >= 5 seconds (limit of temporal resolution) and those of amounts smaller than 5 mg were excluded as such values approximate a limit of detection (non-consummatory contacts are distributed around zero within a ±5 mg range). Imposing this 5-mg threshold resulted in a minimal reduction of total weight consumed per bottle (1.6 mg an average) over the 2-h session and reduced contact bouts/time to a greater degree (a decrease of 22 bouts and 159 seconds), indicating that the threshold largely eliminates non-consumption contacts. Three per-bout measures were recorded – amount, duration and rate. From this high-resolution bout data, drinking behavior within each session was parameterized. These consumption parameters included the following: counts of bouts, sums of bout amounts and durations, maximum bout amount and duration, mean and maximum *within-bout* rate, and latency to 1^st^ bout. All parameters were calculated as per-session measures. Bout counts, sums of bout amounts and durations, and mean *within-bout* rates, were additionally parsed into 40-min time bins collapsed across session to examine within-session changes in consumption.

Measures of liquid consumption amount and rate were normalized to body weight (BW) and ethanol consumption amount and rate were estimated from the combined consumption of 20% and 10% ethanol solutions, adjusted for density of the solutions (20%EtOH*0.166 + 10%EtOH*0.166/2). Mean unit imputation was used to replace missing values due to a software error that caused the loss of data from one session of 3 mice.

Behavioral measures were analyzed with Generalized Linear Mixed Models (GLMM), using SPSS (IBM, v 28). GLMM was chosen as this procedure readily allows for the inclusion of litter as a random effect to control for variance attributable to within-litter (intra-cluster) correlation [91] and greater power than traditional ANOVA-based approaches to within-subject designs [92–94]. Additionally, GLMM (like Generalized Estimating Equations) with SPSS permits choice of distribution (normal, gamma or inverse Gaussian) and link function (defines relationship between linear predictor and outcome; e.g., identity, log, or power functions; eliminates need for, and potential errors of statistical validity from, non-linear transforms of the raw data) [95], Satterthwaite approximation of degrees of freedom (most appropriate for unbalanced designs with complex covariance, reported values are the corrected values), covariance types (Diagonal was used for repeated measures, Variance Components was used for the random effect), and robust estimation of fixed effects and coefficients (to adjust for violations of model assumptions).

Preliminary analyses with GLMM were used to determine the best fit distribution and link function by comparing corrected Akaike Information Criterion (AICc) values of models that only included the intercept and random litter effect (subject block and intercept). A large proportion of data values were zeros and thus a constant was initially added to the raw data to test gamma and inverse Gaussian distributions. However, as the magnitude of this constant was associated with large variations in AICc [96], all final results used only the raw data with a normal distribution and a link function that produced the smallest AICc (the link functions compared were limited to a Tukey’s ladder of powers: values of 2, 1.5, 1[identity], 0.5, 0.25, “0” [log10], −0.25, −0.5, −1, −1.5, −2) excluding those link functions that produced errors. Graphs depict the raw data.

As the detection of interactions was the focus of this study [19,24,47,54] (including replication of prior unpublished observations in our lab that indicated a SCC*GS*LIQ interaction such that XX+**O** drink less ethanol, but not less water, than both XY+**O** and XX+**T**), and as the power to detect interactions are reduced relative to main effects in a full-factorial model [97–100], we also report results from simplified models (e.g., interaction term only) where informative (e.g., effect-size estimates) (See **Supplementary Results - Bout Stats Example**, **Additional File 3**). Importantly, interaction-only models are simply reparameterizations of their respective full-factorial model (same overall model degrees of freedom, AICc and *post hoc* values for sequential Sidak corrected simple effects; lower-level parameters are moved to higher-level interaction terms; thus, higher degrees of freedom and increased power for those higher-level terms).

Significant omnibus effects and planned comparisons were delineated with *post hoc* paired-means comparisons that used the estimated means function of GLMM in SPSS with a sequential Sidak correction for multiple comparisons (*post hoc* p-values represent Sidak corrected values). Cohen’s *f^2^* was used as a measure of effect size for omnibus fixed effects [101] and Cohen’s *d* was used as a measure of effect size in paired-means comparisons; calculated respectively by the *F_to_f2* and *t_to_d* functions of the *effectsize* package of R (The R Foundation for Statistical Computing, version 4.1.3). Conventions for effect size magnitudes of “Large”, “Medium” and “Small” are ≥ 0.35, ≥ 0.15, and ≥ 0.02 for Cohen’s *f*^2^ and ≥ 0.8, ≥ 0.5, and ≥ 0.2 for Cohen’s *d* [102]. Ancillary measures (pre-session body weight, % post-session body weight loss, food consumed, solution preferences, and blood ethanol concentrations; collapsed across session and time bin) were analyzed with bootstrapped ANCOVA with Litter as a random factor due to failures of matrix convergence for Litter with GLMM (example provided in **Supplementary Results - Ancillary Stats Example, Additional File 4**). Correlation tests were 2-tailed non-parametric Spearman’s rho with a false discovery rate correction [103].

## RESULTS

### Effects of Ethanol Concentration on Liquid Consumption Parameters

#### Sums of Bout Amounts and Durations

Liquid (LIQ) consumption (sum of bout *amounts*; g liquid/kg body weight/h) was ~3x greater for **20%** ethanol (M=2.9, SD=2.5) than for either **10%** (M=1.1, SD=1.0) or **0%** (M=0.7, SD=0.7) indicating a strong ethanol preference (LIQ: F_(2,131)_ = 124.9, p < .000001; *f*^2^ = 1.91) (**Figure 1A**). SCC and GS interacted on the consumption of **20%** ethanol (SCC*GS*LIQ: full-factorial model, F_(2,202)_ = 6.9, p = .001, *f*^2^ = 0.07; 3◻way only model, F_(11,197)_ = 57.8, p < .000001, *f*^2^ = 3.23) such that the ranking for **20%** ethanol bout *amounts* was **XX+T** (M=3.9, SD=2.8) **> XY+T** (M=3.5, SD=2.3) **> XY+O** (M=2.7, SD=2.1) **> XX+O** (M=1.6, SD=2.0) (simple effects: XY+**O** > XX+**O**, p = .0498, *d* = 0.48; XX+**T** > XX+**O**, p = .0002, *d* = 1.32; XY+**T** > XY+**O**, p = .037, *d* = 0.32) with no SCC or GS differences in **10%** and **0%** ethanol consumption.

**Figure 1:**
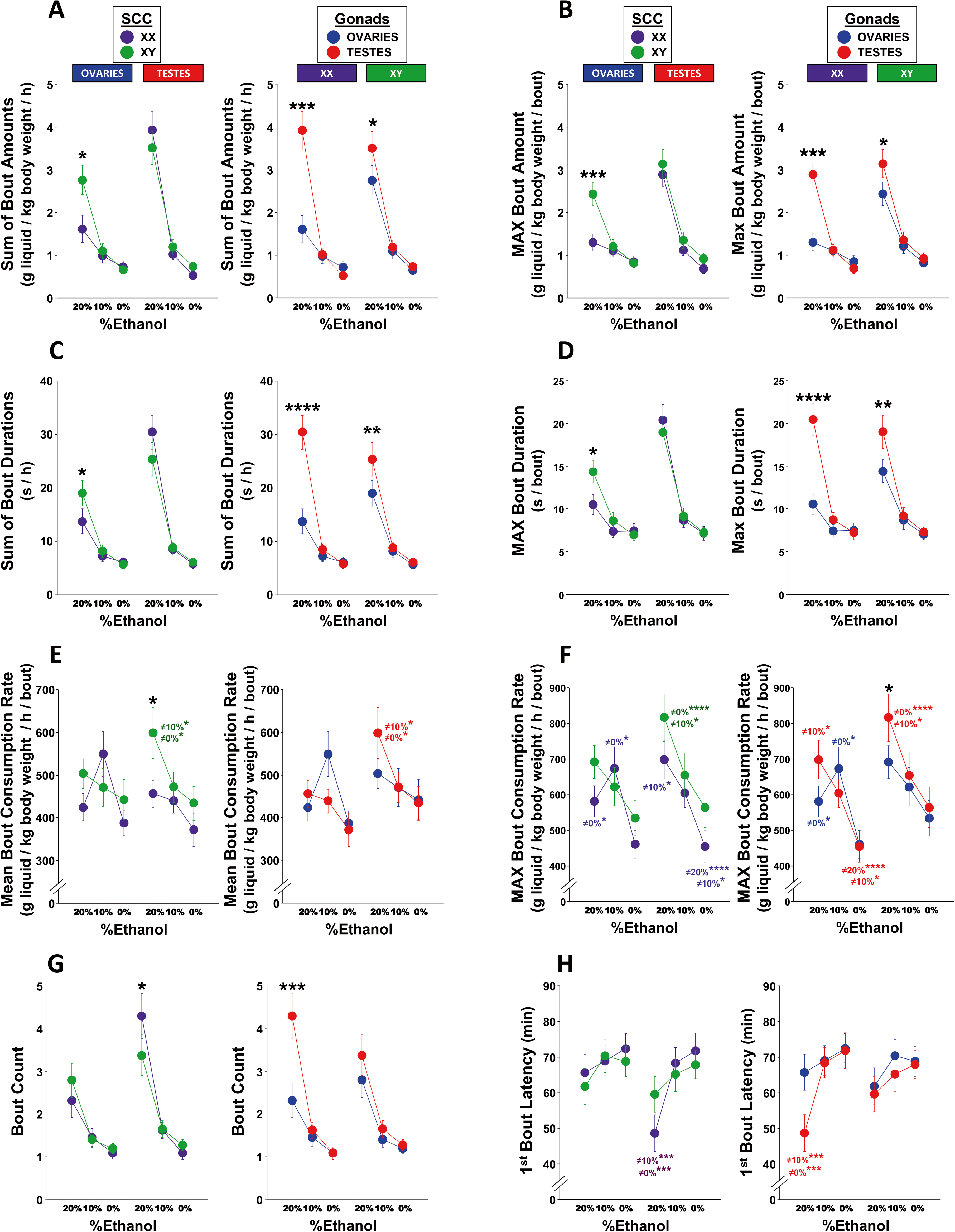
Mean Per-session Bout Parameters as a function of Ethanol Concentration. Figures plot mean values (error bars represent ±SEM) of per-session bout parameters for each ethanol concentration (**20%**, **10%** or **0%**) that are grouped by gonadal sex (**GS**; **OVARIES** or **TESTES**) and split by sex-chromosome complement (**SCC**: **XX** or **XY**) (left side of each letter) or vice versa (right side of each letter) so as to better visualize simple effects of SEX and SCC within levels of the 2^nd^ genotype factor. These parameters include the **Sum of Bout Amounts (A**, g/kg/h), **Maximum (MAX) Bout Amount (B**, g/kg/h/bout), **Sum of Bout Durations (C**, s/h), **Maximum Bout Duration (D**, s/h/bout), **Mean Bout Consumption Rate (E**, g/kg/h/bout; *within-bout rates*), **Maximum Bout Consumption Rate (F**, g/kg/h/bout; *within-bout rate*), **Bout Count (G)**, and **1^st^ Bout Latency (H**, minutes from session start). Sidak adjusted significance values for simple effects of SCC and GS: *p < .05, **p < .01, ***p < .001, ****p < .0001. Sidak adjusted significance values for simple effects of ethanol concentration within genotype use the same asterisk coding for p-values, but are color coded to indicate genotype group and indicate ethanol concentration value from which that mean differs (e.g., ≠**10%\*** in blue represents a significant difference from the 10% ethanol solution in females). For clarity, the simple effects of ethanol concentration within genotype are not included for bout amounts (sum and max), bout durations (sum), and bout counts as all p < .05 except for the following: XX+ovaries 10% vs 0% for sum of bout amounts (p = .1) and durations (p = .2), and for all simple effects in maximum bout amount (all p > .08) and bout duration (all p > .05); other 3 genotypes 10% vs 0% for maximum bout duration (all p > .06). See **Supplementary Results – Liquid** for a full report of statistical analyses and **Supplementary Figure 1** for complementary boxplot-line combo graphs of these data. Group Ns (O=ovaries, T=testes): XX+O=39, XX+T =39, XY+O=38 and XY+T=35.

Similar patterns were observed in the sum of bout *durations* (s/h) (**Figure 1C**): ~3x greater consumption *time* for **20%** (M=22, SD=14) than for either **10%** (M=8, SD=7) or **0%** (M=6, SD=6) (LIQ: F_(2,234)_ = 162.6, p < .000001; *f*^2^ = 1.39) and a **20%** ethanol consumption *duration* ranking of **XX+T** (M=30, SD=20) **> XY+T** (M=25, SD=18) **> XY+O** (M=19, SD=15) **> XX+O** (M=14, SD=15) (SCC*GS*LIQ did not reach statistical significance in the full model [F_(2,283)_ = 2.2, p = .11, *f*^2^ = 0.02], but did in the 3-way only model [F_(11,262)_ = 61.2, p < .000001, *f*^2^ = 2.39]) (simple effects: XY+**O** > XX+**O**, p = .0497, *d* = 0.37; XX+**T** > XX+**O**, p = 0.00003, *d* = 1.17; XY+**T** > XY+**O**, p = .009, *d* = 0.51).

As expected, bout amount and duration were tightly correlated for **20%** ethanol (r_s(151)_ = .95, p < .000001), **10%** ethanol (r_s(151)_ = .90, p < .000001) and **0%** ethanol (r_s(151)_ = .86, p < .000001) (**Supplementary Table T1, Additional File 1**). Thus, as bout amounts were normalized to BW, but bout durations were not, this similar pattern of results indicates that group effects were not strongly dependent upon on group differences in the denominator of the former (i.e., body weight).

#### Bout Counts

Although the total number of drinking bouts per session was low (M=6, SD=4) (**Figure 1G**), the patterns of bout *counts* across ethanol concentrations and between genotypes mirrored that of bout *amounts* and *durations*. A greater *number* of bouts were observed for **20%** ethanol (M=3.2, SD=2.8) than for either **10%** (M=1.5, SD=1.2) or **0%** (M=1.2, SD=0.8) (LIQ: F_(2,183)_ = 94.6; p < .000001; *f*^2^ = 1.03), and **20%** ethanol bout *number* rankings were **XX+T** (M=4.3, SD=3.3) **> XY+T** (M=3.4, SD=2.8) **> XY+O** (M=2.8, SD=2.4) **> XX+O** (M=2.3, SD=2.4) (SCC*GS*LIQ only trended in the full model [F_(2,275)_ = 2.9, p = .056, *f*^2^ = 0.02], but was statistically significant in the 3-way only model [F_(11,202)_ = 55.4, p < .000001, *f*^2^ = 3.02]) (simple effects: XX+**T** > XY+**T**, p = .028, *d* = 0.49; XX+**T** > XX+**O**, p = 0.0008, *d* = 0.87).

#### Mean Bout Rate

Mean consumption rate *within bout* (g liquid/kg BW/h/bout) decreased as ethanol concentration decreased (**20%** M=493, SD=245; **10%** M=483, SD=250; **0%** M=409, SD=236) (LIQ: F_(2,37)_ = 5.4, p = .009; *f*^2^ = 0.29) (**Figure 1E**). However, this pattern varied greatly among genotypes (SCC*GS*LIQ did not reach statistical significance in the full model [F_(2,58)_ = 1.7, p = .19, *f*^2^ = 0.06], but did in the 3-way only model [F_(11,52)_ = 8.5, p < .000001, *f*^2^ = 1.80]). Although XY+**T** mice showed a clear dose-response effect (**20%** > **10%**, p = .03, *d* = 0.47; **20%** > **0%**, p = .03, *d* = 0.52), the other 3 genotypes did not (for simple effects of LIQ all other p > .13). Notably, the mean rate for XX+**O** was *numerically* greater for **10%** [M=549] than either **20%** [M=424] or **0%** [M=387], and for **20%** ethanol the mean was significantly greater for XY+**T** (M=599) than XX+**T** (M=456) (p = 0.02, *d* = 1.06).

Importantly, these seemingly extreme consumption rates (~450 g/kg BW/h/bout) are due to the high resolution of discriminable consummatory bouts. With an average bout duration of ~12 s and bout amount of ~1.5 g/kg BW, the **consumption rate** *during* **a bout is ~80 g** *ethanol***/kg BW/h for the 20% solution**. This microstructure analysis highlights that, in the DID procedure, FCG mice voluntarily consume liquids in a “binge-like” manner (fast rates/bout, a few short bouts per hour) that may approximate the pharmacokinetics of ethanol delivered non-volitionally by low-dose intragastric gavage.

#### 1^st^ Bout Latencies

The average latency (min) of the 1^st^ bout was shorter for **20%** ethanol (M=59, SD=32) than either **10%** (M=68, SD=27) or **0%** (M=70, SD=27) (LIQ: F_(2,223)_ = 11.3, p = .00002; *f*^2^ = 0.10) (**Figure 1H**). The 1^st^ bout latency for **20%** was shorter for XX+**T** (M=49) than for the other three genotypes (XY+**T** M=60, XY+**O** M=62, XX+**O** M=66) (simple effect of GS, XX+**T** < XX+**O**, *p* = 0.07, *d* = 0.73). But, these results should be interpreted with caution as variance was constrained by a 120 min limit (if no bout was observed in a session, the maximum value was assigned; see **Supplementary Figure S1H, Additional File 1**).

#### Maximum Bout Amount, Duration and Rate

For **20%** ethanol, the per-session values for the *maximum single bout amount* (M=2.4 g/kg/bout; **Figure 1B**) and *maximum single bout duration* (M=16 s/bout; **Figure 1D**) were only slightly smaller than the per-hour *sum of bout amounts* (M=2.9 g/kg/h) and *sum of bout durations* (M=22 s/h), indicating that mice drink most of their **20%** ethanol in a single large bout. Moreover, the *maximum within-bout rate* (612 g/kg/h/bout; **Figure 1F**) was quite a bit larger than the *mean within-bout rate* (462 g/kg/h/bout). These observations (and the results below) again indicate binge-like ethanol consumption.

The pattern of ethanol-concentration effects and genotype differences for the *maximum bout amount* (**Figure 1B**) was similar to that of the sum of bout amounts. Larger *maximum* bout *amounts* (g/kg/bout) were observed for **20%** ethanol (M=2.4, SD=1.8) than either **10%** (M=1.2, SD=1.0) or **0%** (M=0.8, SD=0.7) (LIQ: F_(2,123)_ = 70.9, p < .000001; *f*^2^ = 1.15), and the genotype rankings for **20%** ethanol were **XY+T** (M=3.1, SD=2.0) **> XX+T** (M=2.9, SD=1.7) **> XY+O** (M=2.4, SD=1.7) **> XX+O** (M=1.3, SD=1.2) (SCC*GS*LIQ: full model, F_(2,208)_ = 7.3, p = .0008, *f*^2^ = 0.07; 3 way only model, F_(11,209)_ = 59.3, p < .000001, *f*^2^ = 3.12) (simple effects: XY+**O** > XX+**O**, p = .0009, *d* = 0.70; XX+**T** > XX+**O**, p = 0.0001, *d* = 1.04; XY+**T** > XY+**O**, p = .046, *d* = 0.50).

Similarly, the pattern of ethanol-concentration effects and genotype differences for the *maximum bout duration* (**Figure 1D**) was similar to that of the sum of bout durations. Larger *maximum* bout *durations* (s/bout) were observed for **20%** ethanol (M=16, SD=10) than either **10%** (M=8, SD=5) or **0%** (M=7, SD=4) (LIQ: F_(2,182)_ = 68.6, p < .000001; *f*^2^ = 0.75), and the genotype rankings for **20%** ethanol were **XY+T** (M=20, SD=11) ≈ **XX+T** (M=19, SD=11) **> XY+O** (M=14, SD=8) **> XX+O** (M=11, SD=7) (SCC*GS*LIQ did not reach statistical significance in the full model [F_(2,186)_ = 1.1, p = .32, *f*^2^ = 0.01], but did in the 3-way only model [F_(11,211)_ = 22.4, p < .000001, *f*^2^ = 1.71]) (simple effects: XY+**O** > XX+**O**, p = .0009, *d* = 0.70; XX+**T** > XX+**O**, p = 0.0001, *d* = 1.04; XY+**T** > XY+**O**, p = .046, *d* = 0.50).

Lastly, the pattern of the ethanol-concentration effect on *maximum* within-bout *rate* (g liquid/kg BW/h/bout) (**Figure 1F**) was similar to that of the mean bout rate as within-bout *maxima* decreased as ethanol concentration decreased (**20%** M=694, SD=323; **10%** M=638, SD=324; **0%** M=503, SD=287) (LIQ: F_(2,94)_ = 18.0, p < .000001; *f*^2^ = 0.38) while those for the effect of genotype were slightly different. Although XY+**T** again showed a clear dose-response effect (**20%** > **10%**, p = .049, *d* = 0.31; **20%** > **0%**, p = .00002, *d* = 0.44) and XX+**O** again showed greater maxima for **10%** than either **20%** or **0%** (**10%** > **0%**, p = .03, *d* = 0.45; but, **20%** > **0%**, p = .03, *d* = 0.25), XX+**T** also showed a clear dose-response effect (**20%** > **10%**, p = .03, *d* = 0.12; **20%** > **0%**, p = .00005, *d* = 0.25; **10%** > **0%**, p = .03, *d* = 0.21) and for **20%** ethanol, maxima were greater for XY+**T** than XY+**O** (p = .02, *d* = 0.41).

### Patterns of Ethanol Consumption *Within* Session

#### Ethanol Bout Counts, Amounts and Durations

Within-session changes in consumption varied greatly by genotype, with XY+**O** mice consuming more ethanol than XX+**O** mice for only the 1^st^ 80 min, XX+**T** mice consuming more ethanol than XX+**O** mice for the entire session – but, to a greater degree earlier than later – and XY+**T** mice consuming more ethanol the XY+**O** mice only at the beginning and end of sessions (**Figure 2 A-C**).

**Figure 2:**
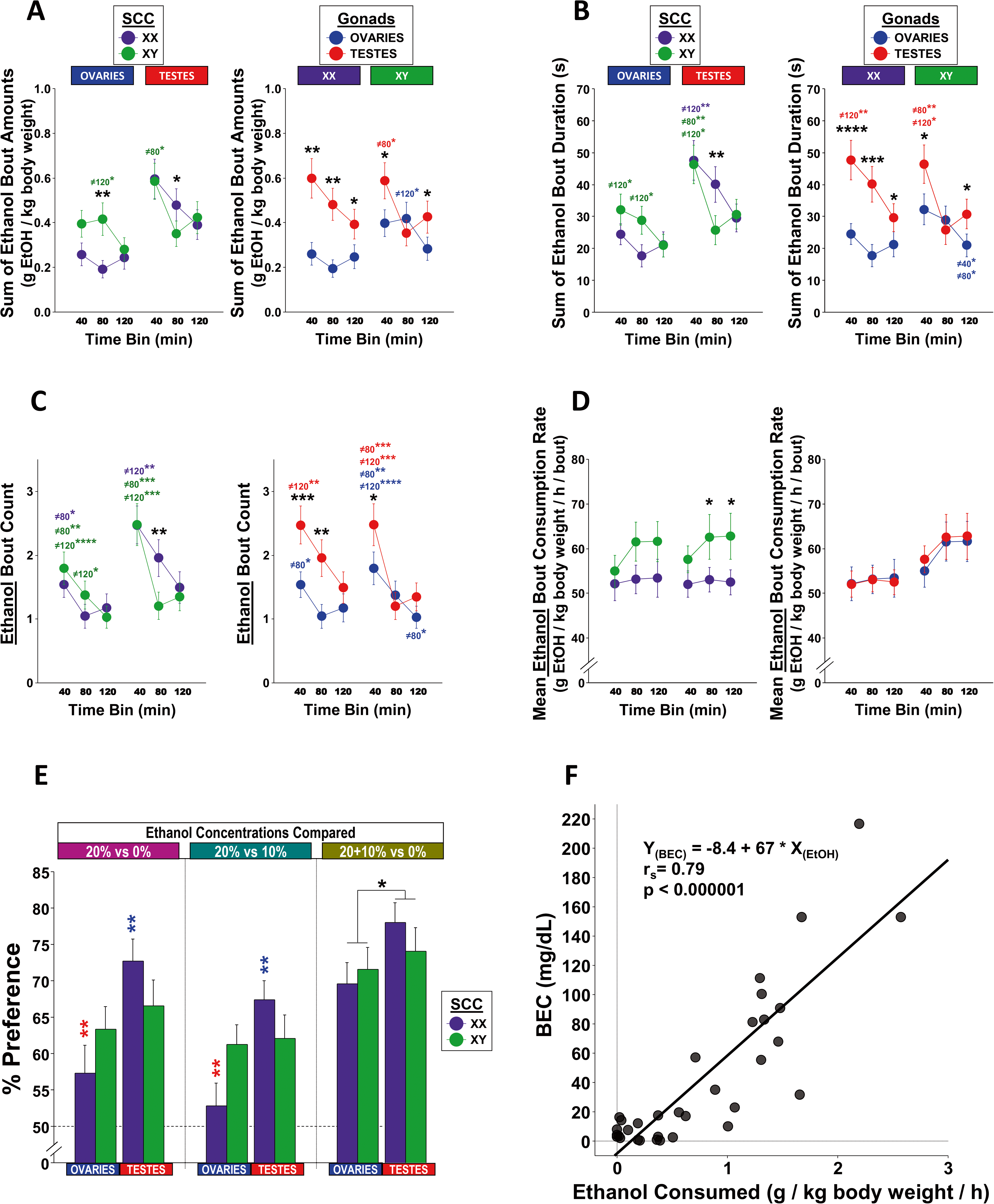
Mean Per-session Ethanol Bout Parameters as a function of Within-Session Time Bin, Blood Ethanol Concentrations (BECs) After Last Session, and Mean Per-session Solution Preferences. **A-D:** Figures plot mean values (error bars represent ±SEM) of time-binned bout parameters for ethanol consumption (**20%** + **10%** ethanol solutions) that are grouped by gonadal sex (**GS**; **OVARIES** or **TESTES**) and split by sex-chromosome complement (**SCC**: **XX** or **XY**) (left side of each letter) or vice versa (right side of each letter) so as to better visualize simple effects of GS and SCC within levels of the 2^nd^ factor. These per-bin parameters include the **Sum of Ethanol-Bout Amounts (A**, g/kg), **Sum of Ethanol-Bout Durations (B**, s), **Count of Ethanol Bouts (C)**, and **Mean Ethanol-Bout Consumption Rate (D**, g/kg/h; *within-bout rates*). Sidak adjusted significance values for simple effects of SCC and GS: *p < .05, **p < .01, ***p < .001, ****p < .0001. Sidak adjusted significance values for simple effects of time bin within genotype use the same asterisk coding for p-values, but are color coded to indicate genotype group and numerically indicate the session from which that session differs (e.g., ≠**40\*** in purple represents a significant difference from the 40-min time bin in XX mice). **E: Preference of Ethanol Concentrations** (%) grouped by the ethanol concentrations compared (dashed line at 50% represents no preference). Sidak adjusted significance values for simple effects of SCC and GS (*p < .05, **p < .01, ***p < .001, ****p < .0001) are color coded to indicate the genotype group from which that mean differs (e.g., **\*\*** in red over the bar for XX mice with ovaries represents a p < .01 significant difference from XX mice with testes. Group Ns for bouts and solution preferences (O=ovaries, T=testes): XX+O=39, XX+T =39, XY+O=38 and XY+T=35. **F: BECs** (mg/dL) as a function of ethanol consumed (EtOH g/kg/h; blood samples were collected within 5 min of the completion of the last session). No statistically significant main effect of either GS or SCC, nor of their interaction, on BECs (ANCOVA, all p > .13), but main effect of EtOH (p < .000001) (Spearman’s rho for BEC vs EtOH: r_s_ (33) = 0.79). Group Ns for BECs: XX+O=11, XX+T=7, XY+O=6, XY+T=9.

GLMM analyses confirmed these patterns for bout *counts* (SCC*GS\***Time Bin** (**TB**; 1 = 1^st^ 40 min, 2 = 2^nd^ 40 min, 3 = last 40 min): full model, F_(2,201)_ = 4.1, p = .018, *f*^2^ = 0.04; 3◻way only model, F_(11,227)_ = 21.3, p < .000001, *f*^2^ = 1.03) (simple effect of **SCC**: XY+**O**_(TB**2**)_ > XY+**O**_(TB**2**)_, p = .007, *d* = 0.62; simple effects of **GS**: XX+**T**_(TB**1**)_ > XX+**O**_(TB**1**)_, p = .0003, *d* = 0.79; XX+**T**_(TB**2**)_ > XX+**O**_(TB**2**)_, p = .004, *d* = 0.74; XY+**T**_(TB**1**)_ > XY+**O**_(TB**1**)_, p = .01, *d* = 0.73; XY+**T**_(TB**3**)_ > XY+**O**_(TB**3**)_, p = .02, *d* = 0.26), bout *amount* sums (SCC*GS*TB: full model, F_(2,159)_ = 4.5, p = .013, *f*^2^ = 0.06; 3◻way only model, F_(11,147)_ = 6.7, p < .000001, *f*^2^ = 0.50) (simple effects of **SCC**: XY+**O**_(TB**2**)_ > XX+**O**_(TB**2**)_, p = .008, *d* = 0.45; XX+**T**_(TB**2**)_ > XY+**T**_(TB**2**)_, p = .046, *d* = 0.46; simple effects of **GS**: XX+**T**_(TB**1**)_ > XX+**O**_(TB**1**)_, p = .001, *d* = 1.17; XX+**T**_(TB**2**)_ > XX+**O**_(TB**2**)_, p = .001, *d* = 0.88; XX+**T**_(TB**3**)_ > XX+**O**_(TB**3**)_, p = .01, *d* = 0.44; XY+**T**_(TB**1**)_ > XY+**O**_(TB**1**)_, p = .04, *d* = 0.63; XY+**T**_(TB**3**)_ > XY+**O**_(TB**3**)_, p = .02, *d* = 0.34), and bout *duration* sums (SCC*GS*TB: full model, F_(2,293)_ = 4.0, p = .02, *f*^2^ = 0.03; 3 way only model, F_(11,292)_ = 14.1, p < .000001, *f*^2^ = 0.53) (simple effects of **SCC**: XY+**O**_(TB**2**)_ > XX+**O**_(TB**2**)_, p = .09, *d* = 0.29; XX+**T**_(TB**2**)_ > XY+**T**_(TB**2**)_, p = .006, *d* = 0.50; simple effects of **GS**: XX+**T**_(TB**1**)_ > XX+**O**_(TB**1**)_, p = .00007, *d* = 0.87; XX+**T**_(TB**2**)_ > XX+**O**_(TB**2**)_, p = .0006, *d* = 0.73; XX+**T**_(TB**3**)_ > XX+**O**_(TB**3**)_, p = .049, *d* = 0.32; XY+**T**_(TB**1**)_ > XY+**O**_(TB**1**)_, p = .01, *d* = 0.54; XY+**T**_(TB**3**)_ > XY+**O**_(TB**3**)_, p = .02, *d* = 0.31).

#### Ethanol Consumption Rate

The *mean* within-bout ethanol consumption *rates* (g/kg/*bin*/bout) of XX mice remained similar across the three 40-min time bins (M=53, SD=16), while consumption rates of XY mice increased over the last 80 minutes (M=62, SD=21) compared to the 1^st^ 40 min (M=56, SD=17), regardless of GS (**Figure 2D**). Although the SCC*TB interaction did not reach statistical significance in either the full model (F_(2,145)_ = 1.6, p = .21, *f*^2^ = 0.02) or 2 way only model (F_(11,169)_ = 1.7, p = .19, *f*^2^ = 0.02), the SCC*GS*TB interaction did reach significance in the 3-way only model (F_(11,138)_ = 2.2, p = .02, *f*^2^ = 0.18) (2-way simple effects of **SCC**: XY_(TB**2**)_ > XX_(TB**2**)_, p = .02, *d* = 0.57; XY_(TB**3**)_ > XX_(TB**3**)_, p = .02, *d* = 0.43; 3-way simple effects of **SCC**: XY+**T**_(TB**2**)_ > XX+**T**_(TB**2**)_, p = .04, *d* = 0.50; XY+**T**_(TB**3**)_ > XX+**T**_(TB**3**)_, p = .04, *d* = 0.45).

### Solution Preferences

At the group level, all genotypes preferred (% ratio of sums of bout amounts) the ethanol solutions (**20%** + **10%**) to water (**0%**) (M=73%, SD=18%); with similar ethanol preferences of **20%** alone over **0%** (M=65%, SD=22%) and **20%** over **10%** (M=61%, SD=19%) (**Figure 2E**). The overall ethanol preference (**20%** +**10%** vs **0%**) was higher in **T** (M=76) than **O** (M=71) (**GS**: F_(1,119)_ = 4.7, p = .03, *f*^2^ = 0.04), regardless of SCC (p > 0.1 for both full and interaction only models). Similarly, preference for **20%** over **0%** was greater in **T** (M=70%) than **O** (M=60%) (**GS**: F_(1,119)_ = 6.3, p = .01, *f*^2^ = 0.05), and preference for **20%** over **10%** was greater in **T** (M=65%) than **O** (M=57%) (**GS**: F_(1,119)_ = 5.1, p = .03, *f*^2^ = 0.04). However, GS and SCC interacted on preference for **20%** over **0%** as XX+**T** (M=73%) showed higher preference than XX+**O** (M=57%) (p = .01, *f*^2^ = 0.06), but no other simple effects (all p > .16) (interaction only model: F_(3,119)_ = 2.8, p = .045, *f*^2^ = 0.07). Similarly, preference for **20%** over **10%** was higher in XX+**T** (M=67%) than XX+**O** (M=53%) (p = .001, *f*^2^ = 0.09), but no other simple effects (all p > .11) (full model: F_(1,119)_ = 5.1, p = .03, *f*^2^ = 0.04; interaction only model: F_(3,119)_ = 3.5, p = .02, *f*^2^ = 0.09).

### Blood Ethanol Concentrations vs Ethanol Consumption

Post-session blood ethanol concentrations (BEC mg/dL: M=42, SD=54, MAX=217, MIN=0; for 8 of 33 mice [24%], BEC ≥80 mg/dL) were positively correlated to within-session ethanol consumption (sum of ethanol bout amounts: g/kg/h) (r_s(33)_ = .79, p < .000001) (**Figure 2F**). ANCOVA confirmed the effect of ethanol consumption on BEC (F_(1,24)_ = 52.9, p < .000001, *f*^2^ = 2.21), but no effects of SCC or GS (all p > .13).

**Patterns of Ethanol Consumption Across Sessions and Ancillary Measures (Pre and Post Session Body Weight and Food Consumption):** See **Supplementary Results - Text, Additional File 2** and **Supplementary Figures and Tables, Additional File 1**.

## DISCUSSION

### Sex-chromosome Complement (SCC) and Gonadal Sex (GS) Interact on Ethanol Binge Drinking in Adolescent Mice

In contrast to many (though not all) ethanol-drinking studies in rodents – especially DID experiments in wildtype mice – for which females (XX with ovaries (**O**)) drink more than males (XY with testes (**T**)) [104,105], our results collected using a modified 3-bottle-choice DID procedure indicate that SCC and GS interact on binge-like ethanol consumption in such a way that mice with testes drink more than mice with ovaries – a difference that is greater in XX compared to XY mice; we also observe that XY mice drink more than XX mice, in the presence of ovaries. These results are more consistent with sex differences in ethanol consumption in humans, for which historically males typically drink more than females across age groups (adults [35-45 y.o.] [106], [18-97 y.o.] [107], young adults [21-30 y.o.] [9], [18-24 y.o.] [20], adolescents [13-17 y.o.] [4], [14-19 y.o.] [8], [13-21 y.o.] [3], and even children [≤12 y.o.] [108]); a difference potentially increasing with developmental age [34,109,110], but decreasing across generations [35–37,111].

Our findings of GS (and SCC) differences in ethanol consumption could be due to one or more procedural differences between our 3 bottle procedure and those using 1-bottle and/or 2◻bottle procedures. These may include the obvious difference – the number of bottles/ethanol concentrations available – and/or the use of adolescent mice (28-31 d.o.) rather than the more common use of adult mice (> 60 d.o.), the use of FCG mice, and the specifics of daily testing (any of the routine procedures involved in daily testing that vary across laboratories).

Like the original Rhodes et al DID procedure [67], many researchers reporting on sex effects on ethanol drinking provide access to only one bottle during a DID session – a bottle of ethanol that replaces the water bottle [67,72,74,75,77–81,84,85]. This 1-bottle procedure is preferred in studies for which achieving high BECs is more important than determining ethanol preferences by way of a ≥2-bottle procedure [73,82,83,112]; Rhodes et al. found higher BECs, but not necessarily higher ethanol intakes, for the 1-over a 2-bottle drinking procedure in 8 inbred mouse strains (including C57BL/6J) [66].

When sex differences were reported in studies using 1-bottle and 2-bottle DID procedures, it was most commonly one wherein females were found to drink more ethanol than males, while differences in preference and/or achieved BECs were less consistent. As ethanol consumption is normalized to body weight, and females weigh less than males, this pattern may indicate that these sex differences in consumption are influenced, in part, by sex differences in body weight. Our results also indicate a modest contribution of body weight, as effect sizes for the GS*SCC interaction on ethanol consumption *amounts* (20%+10% or 20% alone; normalized to body weight) were smaller than that for *durations* (not normalized to body weight). However, not only are GS differences in body weight smaller in adolescence than adulthood, but for the FCG mouse in particular, XX weigh more than XY by PD45 [113,114], a difference that emerges by PD25 [115]. Our results have confirmed this body weight pattern in adolescent FCG mice (PD28-PD31) (see Supplementary Figure S3 A, Additional File 1). Thus, although the SCC effect on ethanol consumption *amounts* in mice with ovaries (XY > XX) is in the same direction as that of the SCC effect on body weight (smaller body weight denominator in XY than XX), this neither explains the GS*SCC interactions on ethanol consumption *amounts* (**T** > **O**; larger effect size in XX than XY mice), nor the similar (though smaller) GS*SCC interactions on *durations*. Thus, as the GS and SCC differences in ethanol consumption reported herein cannot be fully explained by differences in body weight, they more likely reflect group differences in ethanol pharmacokinetics and/or pharmacodynamics.

Interestingly, a 2015 report by Barkley-Levenson and Crabbe, using a line of adult mice selected for high DID BECs (C57BL/6J background), compared 1-bottle to 2-bottle DID procedures and did *not* detect a sex difference in total ethanol consumption using the 1-bottle procedure (but, there were sex differences in bout number – male > female – and bout duration – female > male; indicating different patterns of consumption), but did detect a sex difference using the 2-bottle procedure; the latter was similar to that which we report (males > females) [70]. Additionally, although the study of ethanol drinking in FCG mice by Sneddon et al used 24-h sessions and 5-day testing blocks of escalating ethanol concentration, the effects on ethanol consumption and preference of both GS (**O** > **T**) and SCC (XX>XY) were concentration dependent [24].

Lastly, studies that used a scheduled high alcohol consumption (SHAC) procedures [116] in C57BL/6J mice reported greater ethanol intake in females compared to males in both adolescence and adulthood when either 2-h or 24-h sessions and ethanol solutions from 5% to 20% were used [117], but greater ethanol intake was observed in *adolescent, but not adult*, males compared to females on a version using intermittent 30-min sessions and a 5% ethanol solution [118]. Thus, changes to ethanol concentrations, session scheduling/duration, age of mice and other environmental variables can potentially reverse the anticipated direction of sex differences in ethanol consumption. Considering the importance of resolving challenges to the construct and external validity of preclinical models of voluntary binge-like ethanol consumption like DID (e.g., by determining what factors increase face validity with the ethanol consumption patterns in humans), further studies on the relationship between the aforementioned procedural variations and GS/SCC differences are vital.

However, regardless of the factors that moderate and/or mediate sex differences in models of binge drinking, our microstructural analyses of consummatory behavior further strengthen the construct validity of our procedure. Specifically, most of the ethanol consumption occurred early in the session (i.e., there was “front loading”), was completed in a single, or very few, bouts (bout maxima were only slightly smaller than bout totals), and for the 20% ethanol solution, consumption rate *within a bout* was ~80 g EtOH/kg/h. Moreover, this front-loaded within-session pattern of ethanol consumption was sufficient to maintain intoxicating BEC levels at the end of a final 2-h session in approximately ¼ of the subjects.

### Interaction of GS and SCC on Ethanol Reward and Consumption

Two general mechanisms by which GS and SCC could have interacted on ethanol consumption in our study are: 1) an effect of SCC on pre-adult levels of gonadal hormones and/or responsivity to those hormones (e.g., SCC effects on estrogen and/or androgen receptor expression) or 2) the effects of gonadal hormones depend upon SCC (e.g., sex-chromosomal differences in available response element binding sequences for the nuclear receptors for estrogens and androgens). However, in the FCG mouse, effects of SCC on gonadal hormone levels within GS have yet to be detected either in adulthood [43,119,120] or periadolescence [115]. Thus, although prenatal/perinatal studies have yet to be reported, it seems unlikely that the observed interaction was due to SCC effects on hormone levels in early life.

As mentioned above, human males typically drink more than females across age groups [3,4,8,9,20,106–108] (although this difference potentially increases with age [34,109,110] and appears to be decreasing over generations largely due to increasing alcohol use by females [35–37,111]). One type of environmental change that may impact alcohol consumption in a sex-dependent manner would be changes in exposure to endocrine-disrupting pseudohormones/allohormones (e.g., pseudoestrogens/phytoestrogens from soy products and certain plastics) [89,121] and/or population changes in sex-hormone levels [122,123], as both estrogen (estradiol) and testosterone generally increase alcohol consumption [12–14,17,17,18,124–126]. However, these observations do not help to resolve the relative contributions of, and/or interaction between, the sex-biasing factors of gonadal hormones and SCC.

#### Estrogen and Estrogen Receptors

Evidence indicates that estrogens and SCC can interact in an oppositional manner. Estrogens have been demonstrated to *reduce* body weight and adipose tissue in female mice (Foryst-Ludwig and Kintscher, 2010), while a second X chromosome *increases* these measures (Chen et al., 2012). Moreover, murine studies of cardiovascular ischemia/reperfusion injury have demonstrated a *protective* effect of estrogens (Murphy and Steenbergen, 2014), while a second X chromosome had a *deleterious* effect (Li et al., 2014) (a similar argument can be made for the effects of testosterone and the XY SCC in anxiety – greater when XY than XX, but lower in mice with testes than ovaries [127]). Thus, as estrogens and a 2^nd^ X chromosome can have opposing effects, the loss of either alone may cause an imbalance in XY+**O** and XX+**T** mice. Noting the limitations of DID in terms of ceiling and floor effects in ethanol consumption, we may have been underpowered to consistently detect a difference between XX+**T** and XY+**T** mice (e.g., as shown in Figure 2, XX+**T** mice did consumed more ethanol than XY+**T** mice during the middle 40-min bin of the session).

Additionally, interactions between SCC and gonadal sex on estrogen receptor expression have been reported by Cisternas et al from a study in embryo-derived (E15) amygdala neuronal cell cultures from FCG mice (MF1 mouse genetic background) that detected ~3x greater expression of *Esr1* (estrogen receptor alpha) in XX+**O** neurons, as compared to those from mice of the other 3 genotypes (but, only a main effect of SCC [XY>XX] on *Esr2* [estrogen receptor beta] expression) [128]. Moreover, in a later report they also provide evidence for interactions between SCC and estradiol-dependent axogenesis in embryo-derived (E14) hypothalamic neuron cultures for which neurite outgrowth was greater in neurons from XX+**O** than XY+**O** neurons, and only XY neurons increased neurite growth in response to estradiol (via a Eα receptor mechanism) [129]. Interestingly, although only main effects of SCC on Eα and Eβ receptor mRNA expression were detected (XY > XX), the pattern of results hinted at a SCC*GS interaction on relative Eα expression (XY-XX ≈ 0.8 [p < .05], while XY+**T** -XY+**O** ≈ **0.9**, XY+**T** −XX+**T** ≈ **0.9**, XX+**O** −XX+**T** ≈ **0.15**, and XX+**O** -XY+**O** ≈ **0.15**. Thus, the “differences in differences” interaction was of similar magnitude to the main effect) and Eβ expression (XY-XX ≈ **2.2** [p < .05], while XY+**T** -XY+**O** ≈ **-2.0**, XY+**T** −XX+**T** ≈ **1.6**, XX+**O** −XX+**T** ≈ **0**, and XX+**O** -XY+**O** ≈ **-3.4**). Similarly, a potential 3-way SCC*GS*estradiol interaction on gene expression of neuritogenic factor neurogenin 3 [Ngn3] for which not only did estradiol *increase* expression to a greater degree in XY+**O** than XY+**T**, but estrogen *decreased* expression in XX mice regardless of GS. There was most likely insufficient power in the latter study to detect SCC*GS interactions as it included few subjects (4-6/group) and data were only analyzed with a canonical full-factorial two-way ANOVA [97–100,130,131].

As estrogen (and testosterone) begin to climb toward adult levels around the PD28-PD31 age range used herein [132], and as estrogen increases and/or supports ethanol reward and/or consumption (in mice [126,133–135], in rats [136,137] and in humans [17,138]), greater sensitivity to estrogen in XY mice than XX mice would be consistent with our results in female mice. Moreover, in a human study by de Water et al on a group of Dutch adolescents (11-16 y.o.), although higher estrogen levels were associated with age of alcohol use onset and larger quantities of alcohol consumption in boys, these correlations were not observed in girls (testosterone was also associated with age of use onset in boys, but not girls) [139]. Thus, in both humans and rodents, current evidence indicates that being XY and/or having testes increases sensitivity to alcohol reward enhancement by estrogen (and testosterone: see below), as compared to people who are XX and/or have ovaries.

One potential reason that correlations between gonadal hormones and alcohol consumption are more often detected in males than females may also be that gonadal hormone levels vary over the menstrual cycle. However, current evidence that alcohol consumption is associated with the menstrual cycle is inconsistent [140]. This may be due in part to differences in how the menstrual cycle is measured and whether or not the use of hormone-base contraceptives were included in study designs/models [140,141].

In female wildtype rodents, gonadal hormone levels also fluctuate across the estrous cycle; in C57BL/6J background mice, levels of estradiol are ~7x higher in diestrus and proestrus than in estrous or metestrus [142]. In a study of ethanol consumption using the 1-bottle DID procedure in C57BL/6J mice, differences in ethanol consumption between estrus cycle phases were not detected (nor did DID cause changes in the estrous cycle), even though ovariectomy decreased ethanol consumption and injections of 17β-estradiol benzoate in those ovariectomized increase ethanol consumption [135]. However, in another 1-bottle DID study using C57BL/6J mice, Chen et al compared systemic treatments with selective estrogen receptor degraders that were either high or low in brain penetrance [143]. The high-penetrant compound reduced ethanol consumption in diestrus, but not in estrus (no effects of the low-penetrant compound detected). Interestingly, Chen et al also reported an interaction on estrogen receptor alpha (*Esr1*) expression between DID (ethanol vs water), GS and brain region, such that although ethanol consumption increased *Esr1* expression in the ventral tegmental area of both males and females, it did so in the ventral hippocampus only in males and in the prefrontal cortex only in females. This supports the idea that interactions between sex-biasing factors may more likely be the rule than an exception.

#### Testosterone and Androgen Receptors

The simple effects of GS on ethanol consumption within the GS*SCC interaction that we report herein (mice with ovaries < mice with testes; but to a greater degree in XX mice than XY mice) could also be due in part to increases in ethanol reward caused by testosterone [14]. Levels of testosterone in humans (prenatally exposure; [144], adolescence [12,13,18,139]), and adulthood [144–146]) have been positively associated with ethanol consumption and/or subsequent AUD (but see [147]).

In a report by de Water et al, testosterone levels were associated with age of alcohol use onset in boys, but not girls [139]. Similarly, an earlier study by Costello et al in peripubertal Americans (9-13 y.o.) detected an increased age-corrected likelihood of alcohol use (OR = 1.63) and AUD (OR = 1.95), with increased testosterone levels in boys, but not girls [148]. Additionally, Jones et al reported that the relationship between alcohol use and poorer microstructural integrity in callosal and thalamic white matter in adolescents (~18 y.o.) was explained in part by testosterone (or estradiol), but only in boys [16]. Lastly, Peters et al reported that testosterone increased alcohol intake via reduced amygdala-OFC connectivity, but only in boys [18]. Thus, current evidence indicates that being XY and/or having testes increases sensitivity to alcohol reward enhancement by testosterone (and estrogen; See above) as compared to people who are XX and/or have ovaries.

Additionally, the aforementioned Cisternas et al study in embryo-derived amygdala neuron cultures from FCG mice also indicated a potential interaction between SCC and gonadal sex on androgen receptor expression for which XX+**T** had *greater* expression than XX+**O** (XX+**T** ≈ 2.8, XX+**O** ≈ 1.6; [XX+**T]** −[XX+**O]** ≈ 1.2) while XY+**T** had *lower* expression than XY+**O** (XY+**T** ≈ 1.9, XY+**O** ≈ 2.6; [XY+**T]** −[XY+**O]** ≈ −0.7; thus, an effect size [XX+**T** −XX+**O**] [XY+**T** -XY+**O**] ≈ 1.9 of comparable magnitude to reported main effects) [128]. Moreover, an earlier report by Cisternas et al also indicated potential interactions between SCC and gonadal sex on androgen receptor expression in embryo brains (E16) of FCG mice (MF1 mouse genetic background) that varied by brain region [149]. In the stria terminalis XX+**O** mice had *lower* androgen receptor mRNA expression than the other 3 genotypes (XX+**O** ≈ 0.5, XX+**T** ≈ 1.3, XY+**T** ≈ 1.2, XY+**O** ≈ 1.4; thus, XX+**T** −XX+**O** ≈ **0.8**, XY+**T** -XY+**O** ≈ −0.2, and [XX+**T** −XX+**O**] [XY+**T** -XY+**O**] ≈ **1.0**) and in the anterior amygdala XX+**T** had *greater* expression than the other 3 genotypes (XX+**T** ≈ 1.7, XY+**T** ≈ 0.8, XY+**O** ≈ 0.5, XX+**O** ≈ 0.6; thus, XX+**T** −XX+**O** ≈ **1.1**, XY+**T** -XY+**O** ≈ 0.3, and [XX+**T** −XX+**O**] – [XY+**T** -XY+**O**] ≈ **0.8** – an effect size greater than that reported for the main effect of SCC [XX-XY ≈ **0.45**]).

Lastly, with regard to potential mechanisms for organizational gonadal hormone effects, adult outbred CD-1 mice that were prenatally (E12.5 to E15.5) exposed to androgen receptor inhibition via flutamide showed decreased ethanol consumption compared to vehicle controls, but only in males, while activation by 5α-dihydrotestosterone increased consumption, but only in females [150].

In summary, one explanation for the GS*SCC interaction on ethanol consumption that we report herein is that differences in *testosterone* levels may mediate the higher ethanol consumption in mice with testes compared to mice with ovaries, while an increased sensitivity to *estrogen* (see above) in XY+**O** mice decreases the magnitude of that GS effect within XY mice.

### Genetic Sex-biasing Factors

Ethanol-drinking phenotypes in mice show heritable variation as confirmed by recombinant inbred panel studies [151,152]. This has also been shown for DID in particular by way of selection experiments to generate “high DID” lines of mice via selection based on achieved BECs [70,71,73,80,84,153–156].

Although genetic association studies have identified sex differences in *autosomal* gene variant associations with alcohol use disorder (e.g., *GABRA2* [gamma-aminobutyric acid type A receptor subunit alpha2] [21,157–159] and DRD2 [dopamine receptor D2] [20,160]), the large number of genetic differences that are directly due to variation in SCC remain relatively understudied as the sex chromosomes are typically excluded (esp. Y chromosome) from genome-wide association studies due to statistical challenges like X-chromosome inactivation, X-dosage compensation and the confounding of GS with SCC [165–169]. Despite these challenges, a number of genes located on the X-chromosome have nonetheless been associated with alcohol abuse/use disorder (GWAS supported: MAOA [monoamine oxidase A] [163,168,170], NLGN4X [neuroligin 4 X-linked] [171]; only candidate gene supported: *AR* [androgen receptor] [172–174]).

### X-escapee gene KDM6A

*KDM6A* (X-linked Lysine (K)-specific demethylase 6A or *UTX*) is a H3K27me2/3 demethylase that removes repressive methyl marks on histones to increase expression of genes [175,176]. It escapes X-inactivation in mice and humans and thus is expressed at higher levels in XX relative to XY cells of many tissues, including brain [177–180]. Accordingly, it is a candidate gene for causing sex differences in disease phenotypes that are sensitive to the number of X chromosomes [181–184].

Importantly, GS*SCC interactions on KDM6A (protein) levels in brain homogenates from aged FCG mice (18-22 months) have been reported (XXF ≈ XYF ≥ XXM >XYM) indicating possible regulation of *Kdm6a*/KDM6A by circulating gonadal hormones (e.g., in accord with this pattern, decreased expression caused by androgens; see below) [185]. Moreover, no GS*SCC interaction was detected in hippocampal *Kdm6a* levels in a study on *gonadectomized* adult (3.5-5.5 months) FCG mice (generated by crossing human amyloid protein precursor transgenic [hAPP-J20] female mice with male FCG mice) [182] nor in hypothalamic neuronal cell cultures of FCG mice on the MF1 genetic background [186]. As the experiment with neuronal cultures did not detect an effect of 17β-estradiol on *Kdm6a* levels, the GS*SCC interaction in the experiment using intact mice would more likely be due to androgens.

Genetic association studies have yet to detect a relationship between *KDM6A* and alcohol use, but note the aforementioned bias against inclusion and/or analyses of the X-chromosome in such studies [165–169]. However, in people with early onset AUD (starting use at <21 y.o.), amygdalar brain-derived neurotrophic factor dysregulation was associated with *increased* levels of H3K27me3 (indicating lower H3K27 demethylase activity) [187]. Thus, in XX individuals, the greater relative expression of *KDM6A* may provide some level of protection against the development of AUD by providing a greater buffer against the silencing of this and other K3K27me3-repressed genes silenced by alcohol consumption.

Moreover, *KDM6B* (*autosomal* H3K27me2/3 demethylase) has been shown to be upregulated in AUD and dysregulated in the prefrontal cortex and nucleus accumbens of alcohol dependent rats [188]. Furthermore, ethanol exposure decreases *KDM6B* expression in human dental pulp stem cell [189] and *Kdm6b* expression in rat amygdala [190].

Additionally, in species for which gonadal sex is determined by ambient temperature, H3K27me2/3 demethylation by *KDM6B* promotes activation of the male sex-determining gene *Dmrt1* [191]. Thus, as *SRY* serves the role of sex determination via environmentally *independent* mechanism in humans and mice, it may be the case that *KDM6A* has taken on ontogenic functions both oppositional to *SRY* and with retention of environmental responsivity (e.g., the role of *KDM6A* in responding to hypoxia [192–194]). In other words, *KDM6A* may be an environmentally sensitive feminizing counterforce to *SRY* and the androgenic virilization caused by the effects of *SRY* on gonad differentiation. Moreover, the potential protective function of *KDM6A* against environmental/metabolic challenges [195] may provide a mechanism by which males show greater sensitivity to these challenges, especially in relation to prenatal stressors [196,197] like ethanol exposure [198–201].

### Androgen Receptor (AR) and KDM6A – focus on possible interactions

Both *AR* and *KDM6A* are located on the X-chromosome but are predicted to have opposite effects in terms of driving sex differences in ethanol consumption and AUD (*AR* increasing, *KDM6A* decreasing, see above). However, only *KDM6A* escapes X-inactivation (but, see below). Although the putative contributions of these two factors to sex/gender differences in ethanol consumption are consistent with those observed in humans, they are not consistent with the oft reported sex differences in the rodent literature. Moreover, although we did observe results that were largely consistent with both the human literature and the putative effects of *KDM6A* (XX < XY) and *AR* (**T** > **O**) in that FCG XX+**O** mice drank less ethanol than the other three genotypes (i.e., possible *KDM6A* copy-number effect *sans* AR activation by testicular hormones) and that mice with testes drank more ethanol than those with ovaries (potential effect of AR activation), we also observed a SCC*GS interaction (GS effect greater in XX mice than XY mice and SCC effect only in mice with ovaries). In particular, under these assumptions, XX+**T** mice should have consumed less ethanol that XY+**T** mice (just as XX+**O** mice consume less than XY+**O** mice) as a second X-chromosome would only increase expression of the X-escapee *Kdm6a* – not *Ar*. Furthermore, as discussed above, there is some evidence of tissue-/cell-specific SCC*GS interactions on both *Ar* and *Kdm6a* expression. This raises the possibility that *KDM6A* and *AR* interact via some form of antagonistic regulatory interference (e.g., *KDM6A* downregulates *AR* or vice versa – consistent with the possibility of each driving sex differences in opposite directions) [202].

Although *KDM6A* is an X-escapee gene that has consistently been shown to have greater expression in individuals with two X chromosomes compared to those with only one X chromosome [177–180] and the *AR* gene is typically silenced on the inactive X chromosome [203–205], evidence indicates that *AR* inactivation is not complete (~7.6% allelic imbalance, escape in ~18% of females [206]) and this partial escape varies from one cell type/tissue to another (expression ratio bias overall greater toward females than males in tissues examined; in particular, female bias in pancreas, skin, fibroblasts and spleen, and male bias in tibial nerve and subcutaneous adipose tissue [207]). Similarly, tissue differences in expression levels of both common X-escapee genes like *Kdm6a* and X-genes with significant differences in X-inactivation status have been demonstrated in female *Xist*Δ/+ mice (C57BL/6J background, mice show complete X-inactivation skew of paternal X chromosome) [177].

Some additional evidence in support of *AR*KDM6A* interactions includes studies of castration-resistant (i.e., androgen independent) prostate cancer that implicate loss of H3K27me3 by *KDM6A* and/or *KDM6B* [208]. Moreover, co-immunoprecipitation of AR and KDM6A has been reported in prostate cancer cells (expressing the oncogenic TMPRSS2:ERG gene fusion) [209] and AR has been demonstrated to decrease expression of *KDM6B* [210]. Additionally, the Polycomb repressor complex 2 (PRC2) *adds* H3K27me3 repressive marks [211] and its catalytic subunit (EZH2 [212]) and required subunit for H3K27me3 binding (EED [213]) directly regulate AR binding to hormone response elements [214]. Lastly, querying the protein interaction network between AR, KDM6A, EZH2 and EED as predicted via STRING (a database of known and predicted protein-protein interactions) returns statistically significant protein-protein interaction enrichment in both humans (p = 0.0001) and mice (p = 0.005) [215]. Thus, both direct and indirect interactions between *AR* and *KDM6A* (and/or their gene products) may support their apparent oppositional effects in driving sex differences – masculinization via *AR* [216,217] and feminization via *KDM6A* [202].

### PERSPECTIVES and SIGNIFICANCE

Our results demonstrated that in adolescent mice gonadal sex and sex chromosome complement interact on binge-like ethanol consumption in a pattern (XX+**T** >XY+**T** >XY+**O** >XX+**O**) of greater concordance with human sex differences (**T** > **O**; XY > XX) than those typically reported in rodent models (**O** > **T**; XX > XY). Although further studies would be needed to address the challenge to construct and external validity posed by the discrepancy between the sex differences we observed with our 3-bottle procedure in adolescent mice as compared to that of most other relevant 1-bottle or 2-bottle DID reports typically performed in adult mice (e.g., determine the effects of developmental stage and of the number and range of ethanol concentrations made available on binge-like ethanol consumption), our results not only have greater face validity as a pre-clinical model of binge drinking in adolescent humans but our behavioral microanalysis of the 3-bottle procedure further strengthens the validity of DID as a model of binge drinking in mice on a C57BL/6J background – like the FCG mouse.

## CONCLUSIONS

The combination of our 3-bottle Drinking-in-the-Dark procedure with the use of the FCG mouse may prove to be a very useful approach to examining putative GS by SCC interactions on ethanol consumption that are either X-linked (e.g., putative interactions between *AR* and *KDM6A*) and/or Y-linked (*except* for those involving the *Sry* gene as it remains confounded with GS in the FCG mouse). Results from such lines of investigation may provide valuable insights into the interdependency of genetic and gonadal hormone contributions to generating the sex differences observed in adolescent binge drinking as well as in AUD more generally.

## Supporting information

Additional File 2: Supplementary Results Text

Additional File 3: Supplementary Results Bout Stats Example

Additional File 4: Supplementary Results Ancillary Stats Example

## DECLARATIONS

### Ethics approval and consent to participate

Procedures and protocols were approved by the Institutional Animal Care and Use Committee (IACUC) of the Laboratory Animal Resources division of research at The State of New York (SUNY) Binghamton University

### Consent for publication

Not applicable.

### Availability of data and materials

The datasets generated and analyzed in this study are available from the corresponding author upon reasonable request.

### Authors’ contributions

SMA wrote the manuscript, designed and ran the experiment, and analyzed the data JRB designed and constructed the experimental apparatuses, data capture software, and data transfer scripts

JDJ contributed to the manuscript and design of the experiment, and secured and provided the institutional and financial support needed to run the experiment

All authors read and approved the final manuscript

## Acknowledgements

We thank Barbara Force for her invaluable help with maintaining the FCG breeding colony in our lab and Greg A. Keay Golyakhovsky for technical assistance.

## Additional Files

**Additional File 1 (Supplementary Figures and Tables.pdf):** Includes **Supplementary Figures** (**S1**: Bout parameters as a function of ethanol concentration that additionally show data distributions (box plots), **S2**: Ethanol Bout Parameters Across Sessions, **S3** Ancillary Measures of Initial Body Weight, Post-session Weight Change, Food Consumed) and **Supplementary Tables** (**T1**: Correlations between Bout Parameters and Ancillary Measures, **T2**: Correlations between Blood Ethanol Concentrations, Bout Parameters and Ancillary Measures in a Subset of Mice).

**Additional File 2 (Supplementary Results - Text.docx):** Includes text of results for analyses of ethanol bout parameters as a function of session and for ancillary measures (Initial Body Weight, Post-session Weight Change, Food Consumed).

**Additional File 3 (Supplementary Results – Bout Stats Example.pdf)**: Includes an example SPSS output of GLMM analyses. Included example is the analysis of bout amount sums under various models for the fixed effects of ethanol concentration (liquid type), sex-chromosome complement and gonadal sex (with litter as a random effect).

**Additional File 4 (Supplementary Results – Ancillary Stats Example.pdf)**: Includes example SPSS statistical output of bootstrapped ANOVA analysis of mean pre-session body weight.

