## Additional File 1: Supplementary Figures and Tables for "Gonadal Sex and Sex-Chromosome Complement Interact to Affect Ethanol Consumption in Adolescent Four Core Genotypes Mice"

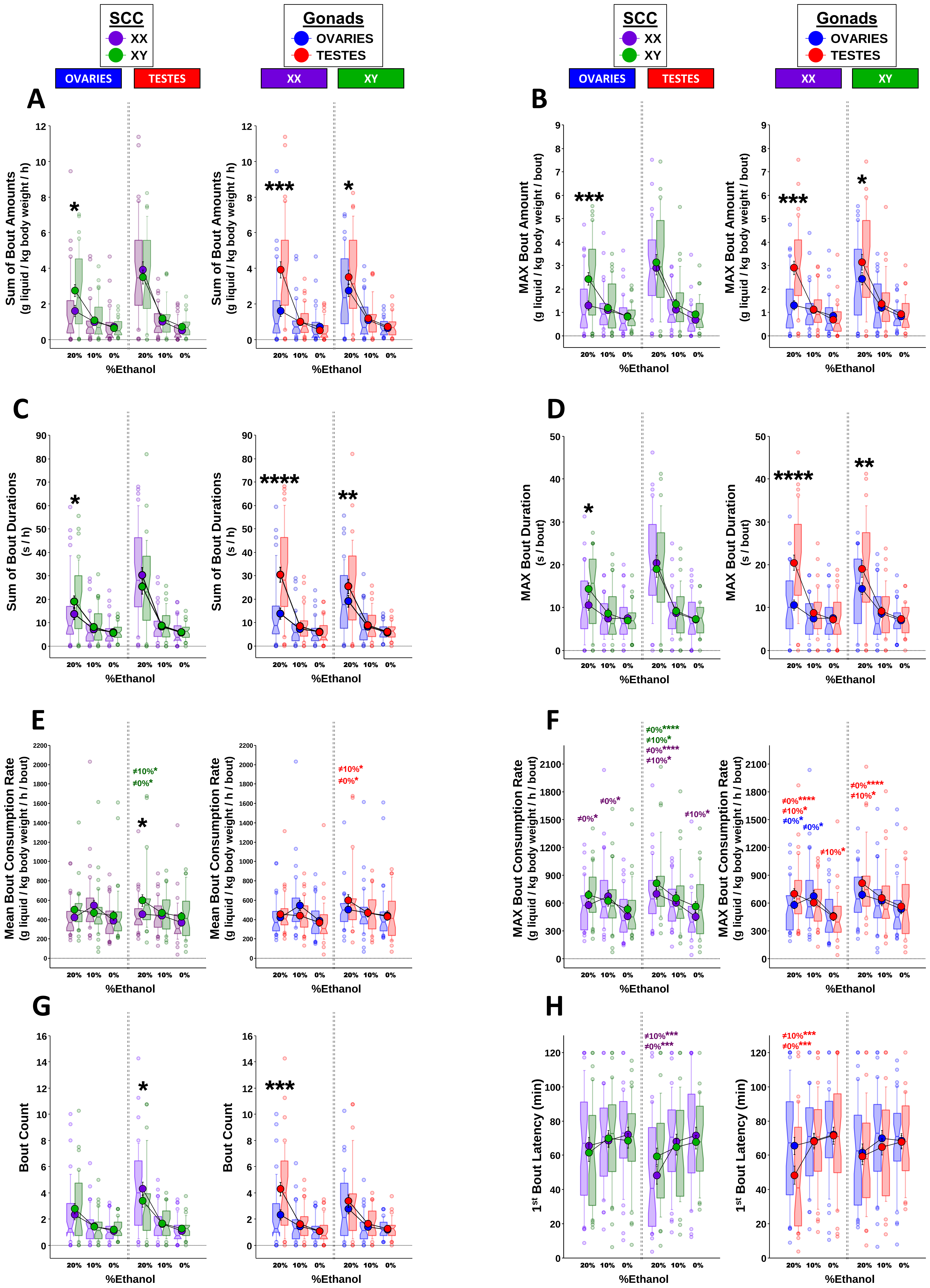

SCC
XX
XY
Gonads
OVARIES
TESTES
OVARIES
TESTES
XX
XY
A
B
C
D
E
F
G
H
Sum of Bout Amounts (g liquid / kg body weight / h)
MAX Bout Amount (g liquid / kg body weight / bout)
Sum of Bout Durations (s / h)
MAX Bout Duration (s / bout)
Mean Bout Consumption Rate (g liquid / kg body weight / h / bout)
MAX Bout Consumption Rate (g liquid / kg body weight / h / bout)
Bout Count
1st Bout Latency (min)
20%
10%
0%
%Ethanol
≠10%*
≠0%*
≠0%****
≠10%*
≠0%****
≠10%*
≠0%*
≠10%*
≠0%****
≠10%*
≠0%*
≠10%*
≠10%***
≠0%***

**Figure S1: Mean Per-session Bout Parameters as a function of ethanol concentration.** Figures are line graphs that depict mean values (larger filled circles, error bars represent ±SEM) overlaid on box-plots (box notch at median; top and bottom of box are $75^{th}$ and $25^{th}$ quartile, respectively; whiskers extend extra $15^{th}$ percentile; small circles represent extreme values) of per-session bout parameters for each ethanol concentration (**20%**, **10%** or **0%**) that are grouped by gonadal sex (**GS**; **OVARIES** or **TESTES**) and split by sex-chromosome complement (**SCC**: **XX** or **XY**) (left side of each pair) or vice versa (right side of each pair) so as to better visualize simple effects of GS and SCC within levels of the $2^{nd}$ factor. These parameters include the **Sum of Bout Amounts (A**, g/kg/h**)**, **Maximum (MAX) Bout Amount (B**, g/kg/h/bout**)**, **Sum of Bout Durations (D**, s/h**)**, **Maximum Bout Duration (E**, s/bout**)**, **Mean Bout Consumption Rate (F**, g/kg/h/bout; *within-bout rates***)**, **Maximum Bout Consumption Rate (G**, g/kg/h/bout; *within-bout rate***)**, **Bout Count (H**, bouts/session**)**, **$1^{st}$ Bout Latency (H**, minutes from session start**)**. Sidak adjusted significance values for simple effects of SCC and GS: *$p < .05$, **$p < .01$, ***$p < .001$, ****$p < .0001$. Sidak adjusted significance values for simple effects of ethanol concentration within genotype use the same asterisk coding for p-values, but are color coded to indicate genotype group and indicate ethanol concentration value from which that mean differs (e.g., **≠10%*** in blue represents a significant difference from the 10% ethanol solution in mice with ovaries). For clarity, the simple effects of ethanol concentration within genotype are not included for bout amounts (sum and max), bout durations (sum), and bout counts as all $p < .05$ except for the following: XX mice with ovaries 10% vs 0% for sum of bout amounts ($p = .1$) and durations ($p = .2$), and for all simple effects in maximum bout amount (all $p > .08$) and bout duration (all $p > .05$); other 3 genotypes 10% vs 0% for maximum bout duration (all $p > .06$). Group Ns (O=ovaries, T=testes): XX+O=39, XX+T =39, XY+O=38 and XY+T=35.

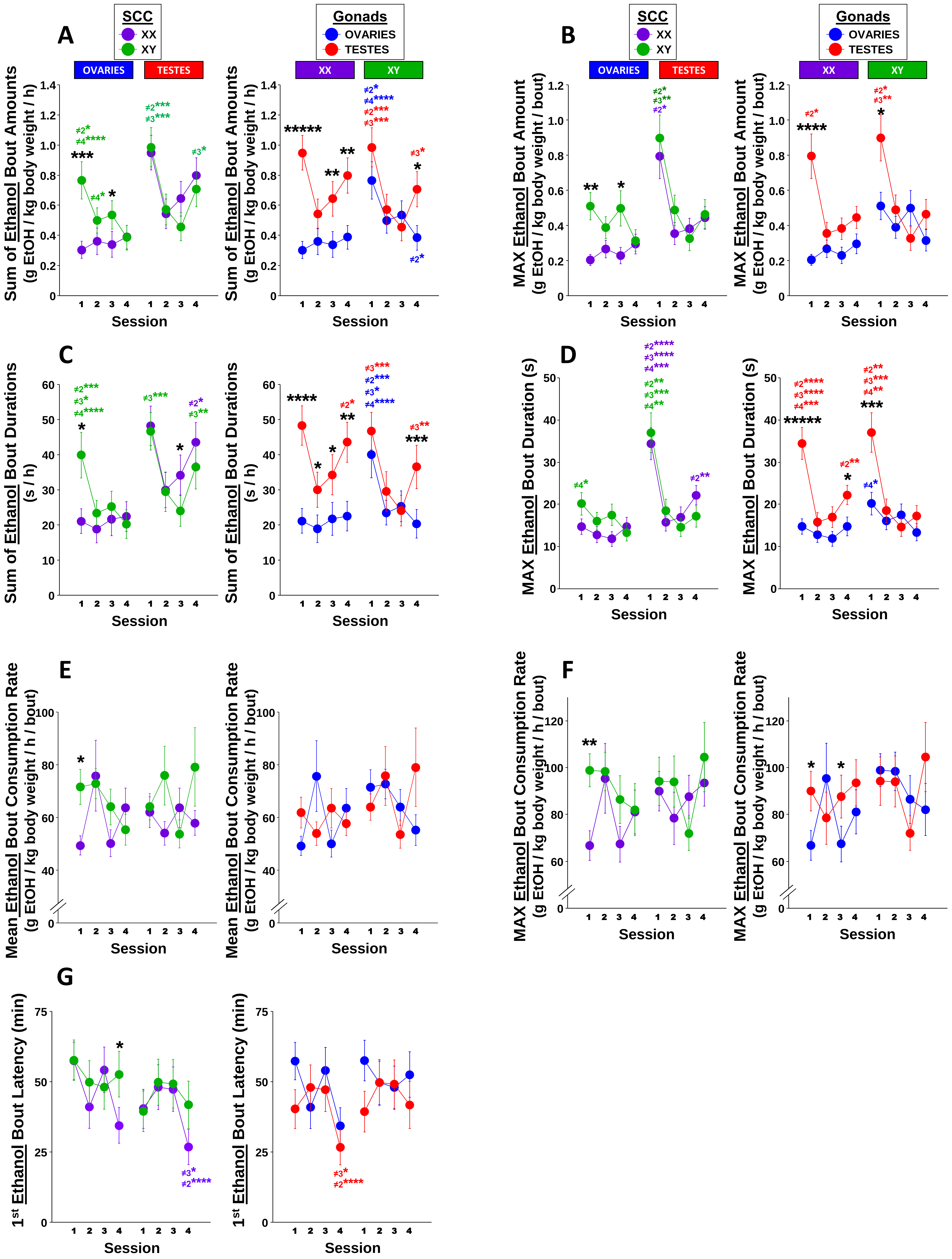
A
B
C
D
E
F
G
SCC
XX
XY
Gonads
OVARIES
TESTES
Sum of Ethanol Bout Amounts
(g EtOH / kg body weight / h)
MAX Ethanol Bout Amount
(g EtOH / kg body weight / bout)
Sum of Ethanol Bout Durations
(s / h)
MAX Ethanol Bout Duration (s)
Mean Ethanol Bout Consumption Rate
(g EtOH / kg body weight / h / bout)
MAX Ethanol Bout Consumption Rate
(g EtOH / kg body weight / h / bout)
1st Ethanol Bout Latency (min)
Session

**Figure S2: Ethanol Bout Parameters as a function of Session.** Figures plot mean values (error bars represent ±SEM) of per-session bout parameters for ethanol consumption (**20%** + **10%** ethanol solutions) that are grouped by gonadal sex (**GS**; **OVARIES** or **TESTES**) and split by sex-chromosome complement (**SCC**: **XX** or **XY**) (left side of each letter) or vice versa (right side of each letter) so as to better visualize simple effects of GS and SCC within levels of the 2nd factor. These parameters include the **Sum of Ethanol-Bout Amounts (A**, g/kg/h**)**, **Maximum (MAX) Ethanol-Bout Amount (B**, g/kg/h/bout**)**, **Sum of Ethanol-Bout Durations (C**, s/h**)**, **Maximum Ethanol-Bout Duration (D**, s/h/bout**)**, **Mean Ethanol-Bout Consumption Rate (E**, g/kg/h/bout; *within-bout rates***)**, **Maximum Ethanol-Bout Consumption Rate (F**, g/kg/h/bout; *within-bout rate***)**, and **1st Ethanol-Bout Latency (G**, minutes from session start**)**. Sidak adjusted significance values for simple effects of SCC and GS: *p < .05, **p < .01, ***p < .001, ****p < .0001. Sidak adjusted significance values for simple effects of session within genotype use the same asterisk coding for p-values, but are color coded to indicate genotype group and numerically indicate the session from which that session differs (e.g., **≠2*** in purple represents a significant difference from the 2nd session in XX mice). Group Ns (O=ovaries, T=testes): XX+O=39, XX+T =39, XY+O=38 and XY+T=35.

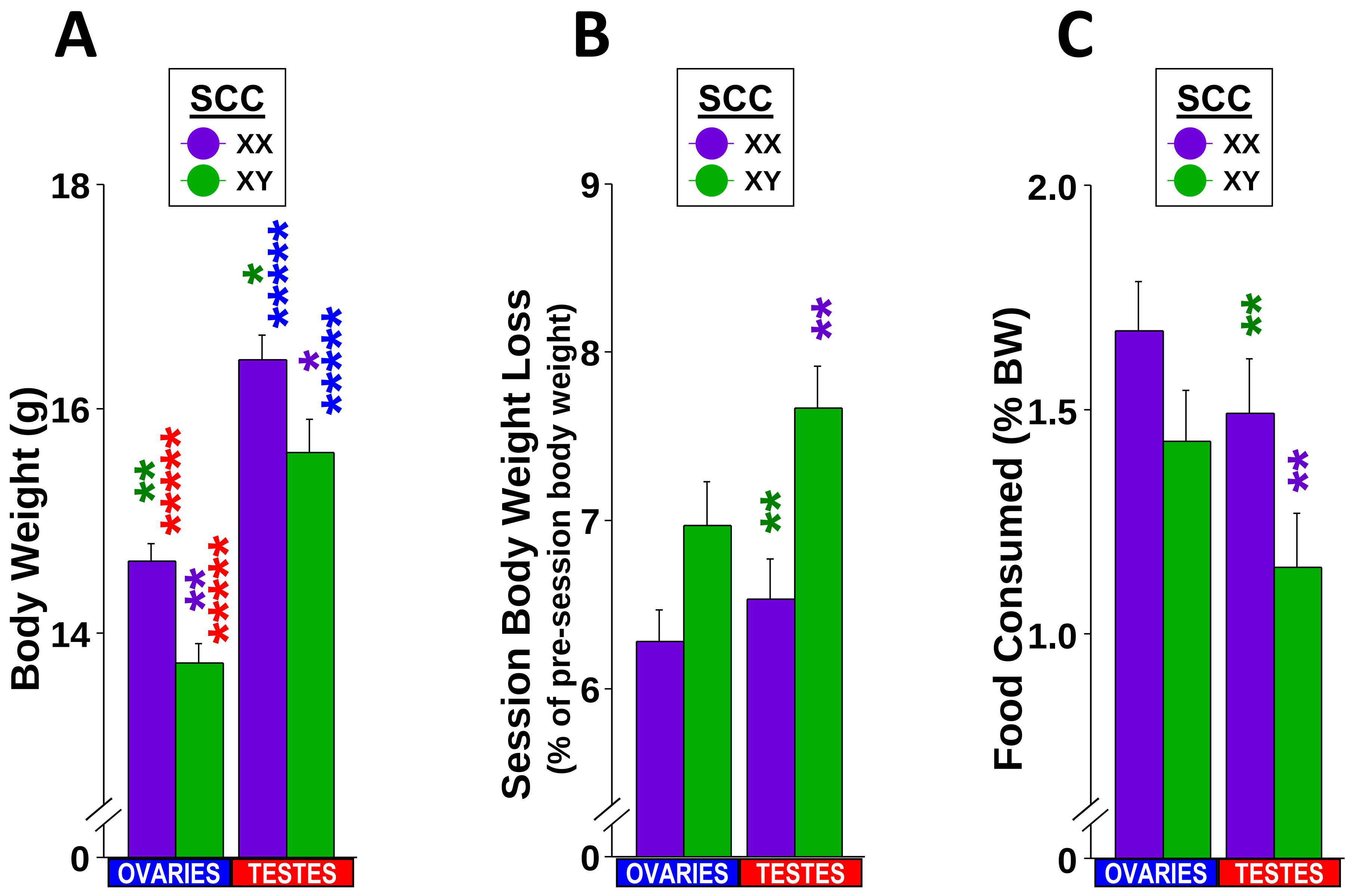

A
SCC
XX
XY
Body Weight (g)
18
16
14
0
OVARIES
TESTES
B
SCC
XX
XY
Session Body Weight Loss
(% of pre-session body weight)
9
8
7
6
0
OVARIES
TESTES
C
SCC
XX
XY
Food Consumed (% BW)
2.0
1.5
1.0
0
OVARIES
TESTES

**Figure S3: Initial Body Weight, Changes in Body Weight, and Food Consumption.** Bars represent mean values (error bars represent ±SEM) of ancillary measures that are grouped by gonadal sex (**GS**; **OVARIES** or **TESTES**) and split by sex-chromosome complement (**SCC**: **XX** or **XY**). These included **Initial Body Weight (A**, g; weight just before 1st Session**)**, **Mean Per-session Body Weight Loss (B**, %100*[post-session weight - pre-session weight]**)**, **Mean Per-session Food Consumed (C**, food consumed during the session as a % of pre-session body weight**)**. Sidak adjusted significance values for simple effects of SCC and GS (*p < .05, **p < .01, ***p < .001, ****p < .0001) are color coded to indicate the genotype group from which that mean differs (e.g., ****** in green over the bar for XX mice with testes represents a p < .01 significant difference from XY mice with testes). Group Ns (O=ovaries, T=testes): XX+O=39, XX+T =39, XY+O=38 and XY+T=35.

### Correlations: 20% ETHANOL

| Spearman's rho (all N = 151) | | AMT SUM | DUR SUM | BOUT COUNT | RATE MEAN | LATENCY | BW PRE | BW LOSS | FOOD |
|---|---|---|---|---|---|---|---|---|---|
| AMT SUM | Correlation Coefficient | 1.000 | 0.952 | 0.910 | 0.473 | -0.740 | 0.080 | -0.049 | 0.334 |
| | Sig. (2-tailed) | | 0.000000 | 0.000000 | 0.000000 | 0.000000 | 0.326794 | 0.553141 | 0.000027 |
| DUR SUM | Correlation Coefficient | 0.952 | 1.000 | 0.960 | 0.260 | -0.774 | 0.142 | -0.052 | 0.297 |
| | Sig. (2-tailed) | 0.000000 | | 0.000000 | 0.001236 | 0.000000 | 0.082423 | 0.524028 | 0.000217 |
| BOUT COUNT | Correlation Coefficient | 0.910 | 0.960 | 1.000 | 0.281 | -0.834 | 0.117 | -0.080 | 0.313 |
| | Sig. (2-tailed) | 0.000000 | 0.000000 | | 0.000470 | 0.000000 | 0.154040 | 0.330744 | 0.000090 |
| RATE MEAN | Correlation Coefficient | 0.473 | 0.260 | 0.281 | 1.000 | -0.285 | -0.112 | -0.044 | 0.244 |
| | Sig. (2-tailed) | 0.000000 | 0.001236 | 0.000470 | | 0.000383 | 0.170677 | 0.588080 | 0.002508 |
| LATENCY | Correlation Coefficient | -0.740 | -0.774 | -0.834 | -0.285 | 1.000 | -0.107 | 0.108 | -0.298 |
| | Sig. (2-tailed) | 0.000000 | 0.000000 | 0.000000 | 0.000383 | | 0.192581 | 0.187594 | 0.000205 |
| BW PRE | Correlation Coefficient | 0.080 | 0.142 | 0.117 | -0.112 | -0.107 | 1.000 | -0.142 | -0.080 |
| | Sig. (2-tailed) | 0.326794 | 0.082423 | 0.154040 | 0.170677 | 0.192581 | | 0.082472 | 0.330088 |
| BW LOSS | Correlation Coefficient | -0.049 | -0.052 | -0.080 | -0.044 | 0.108 | -0.142 | 1.000 | -0.503 |
| | Sig. (2-tailed) | 0.553141 | 0.524028 | 0.330744 | 0.588080 | 0.187594 | 0.082472 | | 0.000000 |
| FOOD | Correlation Coefficient | 0.334 | 0.297 | 0.313 | 0.244 | -0.298 | -0.080 | -0.503 | 1.000 |
| | Sig. (2-tailed) | 0.000027 | 0.000217 | 0.000090 | 0.002508 | 0.000205 | 0.330088 | 0.000000 | |

### Correlations: 10% ETHANOL

| Spearman's rho (all N = 151) | | AMT SUM | DUR SUM | BOUT COUNT | RATE MEAN | LATENCY | BW PRE | BW LOSS | FOOD |
|---|---|---|---|---|---|---|---|---|---|
| AMT SUM | Correlation Coefficient | 1.000 | 0.904 | 0.884 | 0.411 | -0.684 | -0.046 | -0.109 | 0.286 |
| | Sig. (2-tailed) | | 0.000000 | 0.000000 | 0.000000 | 0.000000 | 0.573324 | 0.184022 | 0.000373 |
| DUR SUM | Correlation Coefficient | 0.904 | 1.000 | 0.929 | 0.108 | -0.694 | 0.021 | -0.038 | 0.208 |
| | Sig. (2-tailed) | 0.000000 | | 0.000000 | 0.186868 | 0.000000 | 0.796058 | 0.647507 | 0.010198 |
| BOUT COUNT | Correlation Coefficient | 0.884 | 0.929 | 1.000 | 0.191 | -0.789 | 0.022 | -0.078 | 0.258 |
| | Sig. (2-tailed) | 0.000000 | 0.000000 | | 0.018780 | 0.000000 | 0.790982 | 0.338360 | 0.001372 |
| RATE MEAN | Correlation Coefficient | 0.411 | 0.108 | 0.191 | 1.000 | -0.226 | -0.047 | -0.157 | 0.273 |
| | Sig. (2-tailed) | 0.000000 | 0.186868 | 0.018780 | | 0.005223 | 0.566088 | 0.054159 | 0.000691 |
| LATENCY | Correlation Coefficient | -0.684 | -0.694 | -0.789 | -0.226 | 1.000 | 0.013 | 0.060 | -0.191 |
| | Sig. (2-tailed) | 0.000000 | 0.000000 | 0.000000 | 0.005223 | | 0.876119 | 0.463604 | 0.018774 |
| BW PRE | Correlation Coefficient | -0.046 | 0.021 | 0.022 | -0.047 | 0.013 | 1.000 | -0.142 | -0.080 |
| | Sig. (2-tailed) | 0.573324 | 0.796058 | 0.790982 | 0.566088 | 0.876119 | | 0.082472 | 0.330088 |
| BW LOSS | Correlation Coefficient | -0.109 | -0.038 | -0.078 | -0.157 | 0.060 | -0.142 | 1.000 | -0.503 |
| | Sig. (2-tailed) | 0.184022 | 0.647507 | 0.338360 | 0.054159 | 0.463604 | 0.082472 | | 0.000000 |
| FOOD | Correlation Coefficient | 0.286 | 0.208 | 0.258 | 0.273 | -0.191 | -0.080 | -0.503 | 1.000 |
| | Sig. (2-tailed) | 0.000373 | 0.010198 | 0.001372 | 0.000691 | 0.018774 | 0.330088 | 0.000000 | |

### Correlations: 0% ETHANOL

| Spearman's rho (all N = 151) | | AMT SUM | DUR SUM | BOUT COUNT | RATE MEAN | LATENCY | BW PRE | BW LOSS | FOOD |
|---|---|---|---|---|---|---|---|---|---|
| AMT SUM | Correlation Coefficient | 1.000 | 0.859 | 0.860 | 0.652 | -0.634 | -0.130 | -0.188 | 0.340 |
| | Sig. (2-tailed) | | 0.000000 | 0.000000 | 0.000000 | 0.000000 | 0.110268 | 0.021102 | 0.000020 |
| DUR SUM | Correlation Coefficient | 0.859 | 1.000 | 0.905 | 0.286 | -0.659 | 0.014 | -0.192 | 0.234 |
| | Sig. (2-tailed) | 0.000000 | | 0.000000 | 0.000372 | 0.000000 | 0.860388 | 0.018173 | 0.003873 |
| BOUT COUNT | Correlation Coefficient | 0.860 | 0.905 | 1.000 | 0.361 | -0.781 | -0.050 | -0.137 | 0.268 |
| | Sig. (2-tailed) | 0.000000 | 0.000000 | | 0.000005 | 0.000000 | 0.545967 | 0.092993 | 0.000874 |
| RATE MEAN | Correlation Coefficient | 0.652 | 0.286 | 0.361 | 1.000 | -0.310 | -0.214 | -0.177 | 0.307 |
| | Sig. (2-tailed) | 0.000000 | 0.000372 | 0.000005 | | 0.000106 | 0.008442 | 0.029468 | 0.000126 |
| LATENCY | Correlation Coefficient | -0.634 | -0.659 | -0.781 | -0.310 | 1.000 | -0.033 | 0.150 | -0.162 |
| | Sig. (2-tailed) | 0.000000 | 0.000000 | 0.000000 | 0.000106 | | 0.683026 | 0.065740 | 0.046923 |
| BW PRE | Correlation Coefficient | -0.130 | 0.014 | -0.050 | -0.214 | -0.033 | 1.000 | -0.142 | -0.080 |
| | Sig. (2-tailed) | 0.110268 | 0.860388 | 0.545967 | 0.008442 | 0.683026 | | 0.082472 | 0.330088 |
| BW LOSS | Correlation Coefficient | -0.188 | -0.192 | -0.137 | -0.177 | 0.150 | -0.142 | 1.000 | -0.503 |
| | Sig. (2-tailed) | 0.021102 | 0.018173 | 0.092993 | 0.029468 | 0.065740 | 0.082472 | | 0.000000 |
| FOOD | Correlation Coefficient | 0.340 | 0.234 | 0.268 | 0.307 | -0.162 | -0.080 | -0.503 | 1.000 |
| | Sig. (2-tailed) | 0.000020 | 0.003873 | 0.000874 | 0.000126 | 0.046923 | 0.330088 | 0.000000 | |

**Table T1: Correlations between Bout Parameters and Ancillary Measures.** Table of Spearman's rho correlations between the per-session primary bout measures and ancillary measures split by the three ethanol concentrations (**20%**, **10%** and **0%**). Measures were the sum of bout amounts (AMT SUM), sum of bout durations (DUR SUM), count of bouts (BOUT COUNT), within-bout consumption rate (RATE MEAN), latency to 1$^{st}$ bout (LATENCY), pre-session body weight (BW PRE), post-session body weight loss (BW LOSS), and amount of food consumed during the session (FOOD). Only p-values that survived false discovery rate corrections are highlighted. Note that input values of terms in red font do not vary as a function of ethanol concentration. Group Ns (O=ovaries, T=testes): XX+O=39, XX+T =39, XY+O=38 and XY+T=35.

### Correlations: 20% ETHANOL

| Spearman's rho (all N = 33) | | BEC | AMT SUM | DUR SUM | BOUT COUNT | RATE MEAN | LATENCY | BW PRE | BW LOSS | FOOD |
|---|---|---|---|---|---|---|---|---|---|---|
| BEC | Correlation Coefficient | 1.000 | 0.794 | 0.725 | 0.166 | -0.230 | 0.709 | 0.121 | 0.414 | -0.130 |
| | Sig. (2-tailed) | | 0.000000 | 0.000002 | 0.354489 | 0.196895 | 0.000004 | 0.503123 | 0.016709 | 0.470173 |
| AMT SUM | Correlation Coefficient | 0.794 | 1.000 | 0.944 | 0.262 | -0.443 | 0.891 | -0.018 | 0.492 | -0.113 |
| | Sig. (2-tailed) | 0.000000 | | 0.000000 | 0.140200 | 0.009793 | 0.000000 | 0.921305 | 0.003611 | 0.532579 |
| DUR SUM | Correlation Coefficient | 0.725 | 0.944 | 1.000 | 0.047 | -0.517 | 0.945 | 0.080 | 0.406 | -0.117 |
| | Sig. (2-tailed) | 0.000002 | 0.000000 | | 0.797109 | 0.002043 | 0.000000 | 0.656354 | 0.018901 | 0.516183 |
| BOUT COUNT | Correlation Coefficient | 0.166 | 0.262 | 0.047 | 1.000 | -0.121 | 0.089 | -0.375 | 0.233 | 0.097 |
| | Sig. (2-tailed) | 0.354489 | 0.140200 | 0.797109 | | 0.502410 | 0.623188 | 0.031403 | 0.192009 | 0.590267 |
| RATE MEAN | Correlation Coefficient | -0.230 | -0.443 | -0.517 | -0.121 | 1.000 | -0.641 | -0.005 | -0.012 | -0.140 |
| | Sig. (2-tailed) | 0.196895 | 0.009793 | 0.002043 | 0.502410 | | 0.000059 | 0.979377 | 0.947015 | 0.435882 |
| LATENCY | Correlation Coefficient | 0.709 | 0.891 | 0.945 | 0.089 | -0.641 | 1.000 | 0.116 | 0.326 | -0.020 |
| | Sig. (2-tailed) | 0.000004 | 0.000000 | 0.000000 | 0.623188 | 0.000059 | | 0.521709 | 0.063966 | 0.910853 |
| BW PRE | Correlation Coefficient | 0.121 | -0.018 | 0.080 | -0.375 | -0.005 | 0.116 | 1.000 | -0.342 | 0.046 |
| | Sig. (2-tailed) | 0.503123 | 0.921305 | 0.656354 | 0.031403 | 0.979377 | 0.521709 | | 0.051165 | 0.800913 |
| BW LOSS | Correlation Coefficient | 0.414 | 0.492 | 0.406 | 0.233 | -0.012 | 0.326 | -0.342 | 1.000 | -0.611 |
| | Sig. (2-tailed) | 0.016709 | 0.003611 | 0.018901 | 0.192009 | 0.947015 | 0.063966 | 0.051165 | | 0.000161 |
| FOOD | Correlation Coefficient | -0.130 | -0.113 | -0.117 | 0.097 | -0.140 | -0.020 | 0.046 | -0.611 | 1.000 |
| | Sig. (2-tailed) | 0.470173 | 0.532579 | 0.516183 | 0.590267 | 0.435882 | 0.910853 | 0.800913 | 0.000161 | |

### Correlations: 10% ETHANOL

| Spearman's rho (all N = 33) | | BEC | AMT SUM | DUR SUM | BOUT COUNT | RATE MEAN | LATENCY | BW PRE | BW LOSS | FOOD |
|---|---|---|---|---|---|---|---|---|---|---|
| BEC | Correlation Coefficient | 1.000 | 0.485 | 0.576 | -0.060 | -0.308 | 0.493 | 0.121 | 0.414 | -0.130 |
| | Sig. (2-tailed) | | 0.004205 | 0.000451 | 0.738696 | 0.081130 | 0.003560 | 0.503123 | 0.016709 | 0.470173 |
| AMT SUM | Correlation Coefficient | 0.485 | 1.000 | 0.913 | 0.239 | -0.710 | 0.891 | -0.152 | 0.254 | 0.164 |
| | Sig. (2-tailed) | 0.004205 | | 0.000000 | 0.180696 | 0.000004 | 0.000000 | 0.399620 | 0.153707 | 0.360440 |
| DUR SUM | Correlation Coefficient | 0.576 | 0.913 | 1.000 | -0.086 | -0.707 | 0.939 | -0.085 | 0.318 | 0.069 |
| | Sig. (2-tailed) | 0.000451 | 0.000000 | | 0.632660 | 0.000004 | 0.000000 | 0.639568 | 0.071586 | 0.702939 |
| BOUT COUNT | Correlation Coefficient | -0.060 | 0.239 | -0.086 | 1.000 | -0.065 | -0.022 | -0.176 | -0.136 | 0.348 |
| | Sig. (2-tailed) | 0.738696 | 0.180696 | 0.632660 | | 0.720624 | 0.901971 | 0.327606 | 0.449197 | 0.047451 |
| RATE MEAN | Correlation Coefficient | -0.308 | -0.710 | -0.707 | -0.065 | 1.000 | -0.823 | 0.161 | -0.242 | 0.041 |
| | Sig. (2-tailed) | 0.081130 | 0.000004 | 0.000004 | 0.720624 | | 0.000000 | 0.370297 | 0.175446 | 0.818858 |
| LATENCY | Correlation Coefficient | 0.493 | 0.891 | 0.939 | -0.022 | -0.823 | 1.000 | -0.108 | 0.267 | 0.081 |
| | Sig. (2-tailed) | 0.003560 | 0.000000 | 0.000000 | 0.901971 | 0.000000 | | 0.548465 | 0.133240 | 0.654186 |
| BW PRE | Correlation Coefficient | 0.121 | -0.152 | -0.085 | -0.176 | 0.161 | -0.108 | 1.000 | -0.342 | 0.046 |
| | Sig. (2-tailed) | 0.503123 | 0.399620 | 0.639568 | 0.327606 | 0.370297 | 0.548465 | | 0.051165 | 0.800913 |
| BW LOSS | Correlation Coefficient | 0.414 | 0.254 | 0.318 | -0.136 | -0.242 | 0.267 | -0.342 | 1.000 | -0.611 |
| | Sig. (2-tailed) | 0.016709 | 0.153707 | 0.071586 | 0.449197 | 0.175446 | 0.133240 | 0.051165 | | 0.000161 |
| FOOD | Correlation Coefficient | -0.130 | 0.164 | 0.069 | 0.348 | 0.041 | 0.081 | 0.046 | -0.611 | 1.000 |
| | Sig. (2-tailed) | 0.470173 | 0.360440 | 0.702939 | 0.047451 | 0.818858 | 0.654186 | 0.800913 | 0.000161 | |

### Correlations: 0% ETHANOL

| Spearman's rho (all N = 33) | | BEC | AMT SUM | DUR SUM | BOUT COUNT | RATE MEAN | LATENCY | BW PRE | BW LOSS | FOOD |
|---|---|---|---|---|---|---|---|---|---|---|
| BEC | Correlation Coefficient | 1.000 | 0.219 | 0.168 | 0.085 | -0.106 | 0.354 | 0.121 | 0.414 | -0.130 |
| | Sig. (2-tailed) | | 0.220149 | 0.350120 | 0.639163 | 0.558510 | 0.043516 | 0.503123 | 0.016709 | 0.470173 |
| AMT SUM | Correlation Coefficient | 0.219 | 1.000 | 0.805 | 0.620 | -0.549 | 0.811 | -0.299 | 0.017 | 0.342 |
| | Sig. (2-tailed) | 0.220149 | | 0.000000 | 0.000119 | 0.000935 | 0.000000 | 0.091304 | 0.926457 | 0.051227 |
| DUR SUM | Correlation Coefficient | 0.168 | 0.805 | 1.000 | 0.174 | -0.635 | 0.891 | -0.156 | -0.143 | 0.295 |
| | Sig. (2-tailed) | 0.350120 | 0.000000 | | 0.333135 | 0.000072 | 0.000000 | 0.385244 | 0.428781 | 0.095042 |
| BOUT COUNT | Correlation Coefficient | 0.085 | 0.620 | 0.174 | 1.000 | -0.353 | 0.383 | -0.403 | 0.114 | 0.229 |
| | Sig. (2-tailed) | 0.639163 | 0.000119 | 0.333135 | | 0.043939 | 0.027879 | 0.019936 | 0.527694 | 0.200668 |
| RATE MEAN | Correlation Coefficient | -0.106 | -0.549 | -0.635 | -0.353 | 1.000 | -0.787 | 0.077 | 0.152 | -0.121 |
| | Sig. (2-tailed) | 0.558510 | 0.000935 | 0.000072 | 0.043939 | | 0.000000 | 0.670615 | 0.397155 | 0.503600 |
| LATENCY | Correlation Coefficient | 0.354 | 0.811 | 0.891 | 0.383 | -0.787 | 1.000 | -0.257 | 0.006 | 0.155 |
| | Sig. (2-tailed) | 0.043516 | 0.000000 | 0.000000 | 0.027879 | 0.000000 | | 0.148836 | 0.974038 | 0.390471 |
| BW PRE | Correlation Coefficient | 0.121 | -0.299 | -0.156 | -0.403 | 0.077 | -0.257 | 1.000 | -0.342 | 0.046 |
| | Sig. (2-tailed) | 0.503123 | 0.091304 | 0.385244 | 0.019936 | 0.670615 | 0.148836 | | 0.051165 | 0.800913 |
| BW LOSS | Correlation Coefficient | 0.414 | 0.017 | -0.143 | 0.114 | 0.152 | 0.006 | -0.342 | 1.000 | -0.611 |
| | Sig. (2-tailed) | 0.016709 | 0.926457 | 0.428781 | 0.527694 | 0.397155 | 0.974038 | 0.051165 | | 0.000161 |
| FOOD | Correlation Coefficient | -0.130 | 0.342 | 0.295 | 0.229 | -0.121 | 0.155 | 0.046 | -0.611 | 1.000 |
| | Sig. (2-tailed) | 0.470173 | 0.051227 | 0.095042 | 0.200668 | 0.503600 | 0.390471 | 0.800913 | 0.000161 | |

**Table T2: Correlations between Blood Ethanol Concentrations, Bout Parameters and Ancillary Measures in a Subset of Mice.** Table of Spearman's rho correlations between last-session blood ethanol concentrations, bout measures and ancillary measures for a subset of subjects (N=33) split by the three ethanol concentrations (**20%**, **10%** and **0%**). Measures were the blood ethanol concentrations (BEC), sum of bout amounts (AMT SUM), sum of bout durations (DUR SUM), count of bouts (BOUT COUNT), within-bout consumption rate (RATE MEAN), latency to 1$^{st}$ bout (LATENCY), pre-session body weight (BW PRE), post-session body weight loss (BW LOSS), and amount of food consumed during the session (FOOD). Only p-values that survived false discovery rate corrections are highlighted. Note that input values of terms in red font do not vary as a function of ethanol concentration. No statistically significant main effect of either gonadal sex or sex chromosome complement, nor of their interaction, on BECs (ANCOVA, all $p > .13$), but there was a main effect of total EtOH consumed (sum of bout amounts for **20%**+**10**%) ($p < .000001$). Group Ns (O=ovaries, T=testes): XX+O=11, XX+T=7, XY+O=6, XY+T=9.
