## Additional File 2: Supplementary Results Text for "Gonadal Sex and Sex-Chromosome Complement Interact to Affect Ethanol Consumption in Adolescent Four Core Genotypes Mice"

**Supplemental Results**

Patterns of Ethanol Consumption ***Across*** Sessions:

***Ethanol*** *Bout* ***Amounts*** *and* ***Durations****:* ***Means*** *and* ***Maxima***

Changes across sessions in total per-session consumption of ethanol (i.e., **20%** + **10%** ethanol solutions) varied greatly by genotype (**Supplementary Figure S2 A-D, Additional File 1**). Despite being ethanol naïve, the greatest ethanol consumption ***amounts*** (***sums***, M=0.575 g/kg/h; ***maxima***, M=0.424 g/kg/bout) and ***durations*** (***sums***, M=30.3 s/h; ***maxima***, M=18.5 s/bout) were observed on the 1^st^ session for all genotypes except that of XX+**O** mice. However, although consumption in XX+**O** mice started relatively low and remained low across all four sessions, consumption by mice of the other three genotypes dropped on the 2^nd^ session (~50%). For XY+**O** mice, consumption continued to drop until it reached levels equivalent to that of XX+**O** mice while both XX+**T** and XY+**T** increased to near initial levels over the remaining two sessions. These data indicate that GS and SCC interact on both initial responsivity to ethanol (XX+**O** being less responsive than the other 3 genotypes) and to changes in ethanol consumption over repeated days of DID (XY+**O** continue to decrease while mice with testes of either SCC increase after the 2^nd^-session dip).

Analysis of ethanol consumption ***amounts*** confirmed this pattern across sessions (S) for both ***sums*** (SCC*GS*S: full model, F_(3,122)_ = 2.375, p = .07, *f* ^2^ = 0.06; 3‑way only model, F_(15,186)_ = 9.0, p < .000001, *f* ^2^ = 0.73) (simple effects of **SCC**: XY+**O**_(S_**_1_**_)_ > XX+**O**_(S_**_1_**_)_, p = .0007, *d* = 0.60; XX+**T**_(S_**_3_**_)_ > XY+**T**_(S_**_3_**_)_, p = 0.04, *d* = 0.73; simple effects of **GS**: XX+**T**_(S_**_1_**_)_ > XX+**O**_(S_**_1_**_)_, p < .000001, *d* = 1.03; XX+**T**_(S_**_3_**_)_ > XX+**O**_(S_**_3_**_)_, p = .009, *d* = 0.87; XX+**T**_(S_**_4_**_)_ > XX+**O**_(S_**_4_**_)_, p = .006, *d* = 1.00; XY+**T**_(S_**_1_**_)_ > XY+**O**_(S_**_1_**_)_, p = .007, *d* = 0.47; XY+**T**_(S_**_4_**_)_ > XY+**O**_(S_**_4_**_)_, p = .01, *d* = 0.48) and ***maxima*** (SCC*GS*S: full model, F_(3,40)_ = 3.5, p = .02, *f* ^2^ = 0.26; 3‑way only model, F_(15,120)_ = 8.3, p < .000001, *f* ^2^ = 1.04) (simple effects of **SCC**: XY+**O**_(S_**_1_**_)_ > XX+**O**_(S_**_1_**_)_, p = .001, *d* = 0.37; XY+**O**_(S_**_3_**_)_ > XX+**O**_(S_**_3_**_)_, p = .01, *d* = 0.83; simple effects of **GS**: XX+**T**_(S_**_1_**_)_ > XX+**O**_(S_**_1_**_)_, p = .00001, *d* = 1.63; XY+**T**_(S_**_1_**_)_ > XY+**O**_(S_**_1_**_)_, p = .01, *d* = 1.06).

Lastly, analysis of ethanol consumption ***durations*** also largely confirmed this pattern across sessions for both ***sums*** (SCC*GS*S: full model, F_(3,216)_ = 1.05, p = .37, *f* ^2^ = 0.01; 3‑way only model, F_(15,289)_ = 9.9, p < .000001, *f* ^2^ = 0.51) (simple effects of **SCC**: XY+**O**_(S_**_1_**_)_ > XX+**O**_(S_**_1_**_)_, p = .02, *d* = 0.39; XX+**T**_(S_**_3_**_)_ > XY+**T**_(S_**_3_**_)_, p = 0.03, *d* = 0.54; simple effects of **GS**: XX+**T**_(S_**_1_**_)_ > XX+**O**_(S_**_1_**_)_, p = .00002, *d* = 0.66; XX+**T**_(S_**_2_**_)_ > XX+**O**_(S_**_2_**_)_, p = .02, *d* = 0.50; XX+**T**_(S_**_3_**_)_ > XX+**O**_(S_**_3_**_)_, p = .04, *d* = 0.61; XX+**T**_(S_**_4_**_)_ > XX+**O**_(S_**_4_**_)_, p = .001, *d* = 0.75; XY+**T**_(S_**_4_**_)_ > XY+**O**_(S_**_4_**_)_, p = .0005, *d* = 0.48) and ***maxima*** (SCC*GS*S: full model, F_(3,112)_ = 1.1, p = .36, *f* ^2^ = 0.03; 3‑way only model, F_(15,221)_ = 9.9, p < .000001, *f* ^2^ = 0.67) (simple effects of **SCC**: XX+**T**_(S_**_4_**_)_ > XY+**T**_(S_**_4_**_)_, p = .07, *d* = 0.53; simple effects of **GS**: XX+**T**_(S_**_1_**_)_ > XX+**O**_(S_**_1_**_)_, p < .000001, *d* = 0.86; XX+**T**_(S_**_4_**_)_ > XX+**O**_(S_**_4_**_)_, p = .04, *d* = 0.63; XY+**T**_(S_**_1_**_)_ > XY+**O**_(S_**_1_**_)_, p = .0001, *d* = 1.16).

***Mean*** *and* ***Maximum*** *Ethanol Within-Bout* ***Rate****:*

The ***mean*** and ***maximum*** within-bout ethanol consumption rates across genotypes did not clearly trend upward or downward across sessions, but large fluctuations in rates across sessions were observed both within genotypes and between genotypes (**Supplementary Figure S2 E-F, Additional File 1**). These swings in ethanol consumption rates were confirmed by analysis of both within-bout rate ***means*** (SCC*GS*S: full model, F_(3,41)_ = 7.7, p = .0003, *f* ^2^ = 0.57; 3‑way only model, F_(15,36)_ = 4.3, p < .0002, *f* ^2^ = 1.80) (simple effects of **SCC**: XY+**O**_(S_**_1_**_)_ > XX+**O**_(S_**_1_**_)_, p = .02, *d* = 0.88; XY+**T**_(S_**_2_**_)_ > XX+**T**_(S_**_2_**_)_, p = .07, *d* = 0.59; simple effects trends for **GS**: XY+**T**_(S_**_1_**_)_ > XY+**O**_(S_**_1_**_)_, p = .08, *d* = 0.37; XY+**T**_(S_**_3_**_)_ > XY+**O**_(S_**_3_**_)_, p = .09, *d* = 0.48) and ***maxima*** (SCC*GS*S: full model, F_(3,81)_ = 2.6, p = .06, *f* ^2^ = 0.10; 3‑way only model, F_(15,99)_ = 2.6, p = .002, *f* ^2^ = 0.40) (simple effects of **SCC**: XY+**O**_(S_**_1_**_)_ > XX+**O**_(S_**_1_**_)_, p = .002, *d* = 0.46; XY+**O**_(S_**_3_**_)_ > XX+**O**_(S_**_3_**_)_, p = .08, *d* = 0.54; simple effects for **GS**: XX+**T**_(S_**_1_**_)_ > XX+**O**_(S_**_1_**_)_, p = .03, *d* = 0.42; XX+**T**_(S_**_3_**_)_ > XX+**O**_(S_**_3_**_)_, p = .03, *d* = 0.47).

*1^st^ Ethanol Bout* ***Latencies****:*

Latency (min) to 1^st^ ethanol bout (**20%** or **10%** solution) was stable across the 1^st^ three sessions (S) (S1, M=49, SD=44; S2, M=47, SD=48; S3, M=50, SD=49) then dropped on the 4^th^ session (M=39, SD=45) – an effect driven mostly by XX+**T** (M=27) and XX+**O** (M=34) groups (**Supplementary Figure S2G, Additional File 1**). GLMM analysis confirmed this SCC*S interaction (full model, F_(3,198)_ = 2.7, p = .046, *f* ^2^ = 0.04; 2‑way only model, F_(12,252)_ = 2.7, p = .048, *f* ^2^ = 0.03) (simple effect of **SCC**: XX_(S_**_4_**_)_ < XY_(S_**_4_**_)_, p = .001, *d* = 0.68).

Ancillary Measures

*Pre and Post Session Body Weight and Food Consumption:*

Mice were weighed daily ~30 min before the start of the dark phase and again after the completion of each DID session. The amount of standard chow consumed during these sessions was also recorded. Consistent with prior reports in adolescent FCG mice (Chen et al., 2013), pre-session body weights (g) were larger in **T** (M=16.0, SD=1.6) than **O** (M=14.2, SD=1.1) and larger in XX mice (M=15.5, SD=1.5) than XY mice (M=14.6, SD=1.7) (**GS**: F_(1,119)_ = 76.8, p < .000001, *f* ^2^ = 0.65; **SCC**: F_(1,119)_ = 12.2, p = .0007, *f* ^2^ = 0.10) (**Supplementary Figure S3A, Additional File 1**). Pre-session body weight did not significantly correlate with other ancillary measures nor basic bout measures (sums of bout amounts, sums of bout duration, mean within-bout rate, 1^st^-bout latency) for **20%**, **10%** or **0%** ethanol except for a negative correlation with mean within-bout rate for **0%** (r_s(151)_ = -.21, p = .008) (**Supplementary Table T1, Additional File 1)**.

Most mice lost weight (M=6.8%, SD=1.5%) by the end of the session (**Supplementary Figure S3B, Additional File 1**) and these differences varied by SCC with a larger loss in XY mice (M=7.3%) than XX mice (M=6.4%) (**SCC**: F_(1,119)_ = 11.5, p = .0009, *f* ^2^ = 0.10; but, SCC*GS only model: F_(3,119)_ = 5.1, p = .002, *f* ^2^ = 0.13; thus, likely an attenuated interaction (Blake and Gangestad, 2020)). As expected, weight loss was negatively correlated with food consumed (r_s(151)_ = -.50, p < .000001) (**Supplementary Table T1, Additional File 1)**. However, like pre-session weight, weight loss was not correlated with basic bout measures for **20%** and **10%** ethanol solutions, but for **0%** ethanol weight loss was negatively correlated with bout amounts (r_s(151)_ = -.19, p = .02), bout durations (r_s(151)_ = -.19, p = .02), and within-bout rate (r_s(151)_ = -.18, p = .03).

Lastly, SCC also affected the amount of food consumed (% of pre-session body weight) as consumption was higher in XX mice (M=1.6%, SD=0.7%) than XY mice (M=1.3%, SD=0.7%) (**SCC**: F_(1,119)_ = 9.2, p = .002, *f* ^2^ = 0.08; but, SCC*GS only model: F_(3,119)_ = 4.3, p = .006, *f* ^2^ = 0.11; thus, again, likely an attenuated interaction) (**Supplementary Figure S3C, Additional File 1**). However, unlike body weight and body weight changes, food consumed was *positively* correlated with bout measures of count (**20%**: r_s(151)_ = .31, p = .00009; **10%**: r_s(151)_ = .26, p = .001; **0%**: r_s(151)_ = .27, p = .0008), amount (**20%**: r_s(151)_ = .33, p = .00003; **10%**: r_s(151)_ = .29, p = .0004; **0%**: r_s(151)_ = .34, p = .0002), duration (**20%**: r_s(151)_ = .30, p = .0002; **10%**: r_s(151)_ = .21, p = .01; **0%**: r_s(151)_ = .23, p = .004) and rate (**20%**: r_s(151)_ = .24, p = .003; **10%**: r_s(151)_ = .27, p = .0007; **0%**: r_s(151)_ = .31, p = .0001), and *negatively* correlated with latency (**20%**: r_s(151)_ = -.30, p = .0002; **10%**: r_s(151)_ = -.19, p = .019; **0%**: r_s(151)_ = ‑.16, p < .05 [n.s.]) (**Supplementary Table T1, Additional File 1**).

Thus, overall, although drinking behavior was coupled to food consumption, it was largely decoupled from body weight or body weight changes. This, along with the analyses above, again indicate that the effects of genotype on weight normalized ethanol consumption (20% ethanol in particular) are not directly attributable to genotype differences in weight (especially, for the simple effect of GS in XX mice only on 20% ethanol consumption as the direction of effect for the body weight confound would be in the opposite direction of that observed).
