## Additional File 3: Supplementary Results Bout Stats Example for "Gonadal Sex and Sex-Chromosome Complement Interact to Affect Ethanol Consumption in Adolescent Four Core Genotypes Mice"

**Additional File 3 (Supplementary Results – Bout Stats Example.pdf):**

**Example SPSS output of GLMM analysis of the "sum of bout amounts" parameter under various models for the fixed effects of ethanol concentration (liquid type), sex-chromosome complement and gonadal sex (with litter as a random effect)**

| KEY | | |
|---|---|---|
| SCC | = | Sex Chromosome Complement |
| GT | = | Gonadal Sex |
| LIQ | = | Liquid (20%, 10% or 0% ethanol solution) |
| O | = | Ovaries |
| T | = | Testes |
| 1 (LIQ) | = | 20% EtOH |
| 2 (LIQ) | = | 10% EtOH |
| 3 (LIQ) | = | 0% EtOH |
| AMT_SUM | = | Sum of bout amounts (g liquid / kg body weight / h) |
| GLMM | = | Generalized Linear Mixed Models |
| PH | = | post hoc paired-means comparisons |
| d | = | Cohen's *d* |
| $f^2$ | = | Cohen's *f* |

**FULL FACTORIAL**

**Model Summary**

| | | |
|---|---|---|
| Target | | AMT_SUM |
| Probability Distribution | | Normal |
| Link Function | | Power(.25) |
| Information Criterion | Akaike Corrected | 57.054 |
| | Bayesian | 73.318 |

Information criteria are based on the -2 log likelihood (48.962) and are used to compare models. Models with smaller information criterion values fit better.

**Fixed Effects[a]**

| Source | F | df1 | df2 | Sig. | $f^2$ | 95% CI $f^2$ | | size |
|---|---|---|---|---|---|---|---|---|
| Corrected Model | 57.818 | 11 | 208 | 0.000000000 | 3.06 | 2.30 | 3.81 | H |
| **GS** | **5.688** | **1** | **77** | **0.019540351** | 0.07 | 0.00 | 0.25 | S |
| SCC | 0.581 | 1 | 87 | 0.447809114 | | | | |
| **LIQUID** | **124.910** | **2** | **131** | **0.000000000** | 1.91 | 1.29 | 2.61 | H |
| GS * SCC | 0.222 | 1 | 55 | 0.639632291 | | | | |
| **GS * LIQUID** | **11.015** | **2** | **205** | **0.000028607** | 0.11 | 0.03 | 0.21 | S |
| SCC * LIQUID | 0.263 | 2 | 186 | 0.768919145 | | | | |
| **GS * SCC * LIQUID** | **6.899** | **2** | **202** | **0.001264849** | 0.07 | 0.01 | 0.15 | S |

Probability distribution: Normal Link function: Power(.25)

a. Target: AMT_SUM

**MAIN EFFECTS ONLY**

**Model Summary**

| | | |
|---|---|---|
| Target | | AMT_SUM |
| Probability Distribution | | Normal |
| Link Function | | Power(.25) |
| Information Criterion | Akaike Corrected | 39.101 |
| | Bayesian | 55.430 |

Information criteria are based on the -2 log likelihood (31.011) and are used to compare models. Models with smaller information criterion values fit better.

**Fixed Effects[a]**

| Source | F | df1 | df2 | Sig. | $f^2$ | 95% CI $f^2$ | | size |
|---|---|---|---|---|---|---|---|---|
| Corrected Model | 60.682 | 4 | 227 | 0.000000000 | 1.07 | 0.75 | 1.41 | H |
| SCC | 0.392 | 1 | 168 | 0.532244697 | | | | |
| **GS** | **3.971** | **1** | **131** | **0.048380217** | 0.03 | 0.00 | 0.12 | S |
| **LIQUID** | **115.426** | **2** | **246** | **0.000000000** | 0.94 | 0.66 | 1.25 | H |

Probability distribution: Normal Link function: Power(.25)

a. Target: AMT_SUM

**Up to 2-WAY ONLY**

**Model Summary**

| | | |
|---|---|---|
| Target | | AMT_SUM |
| Probability Distribution | | Normal |
| Link Function | | Power(.25) |
| Information Criterion | Akaike Corrected | 51.770 |
| | Bayesian | 68.053 |

Information criteria are based on the -2 log likelihood (43.679) and are used to compare models. Models with smaller information criterion values fit better.

**Fixed Effects[a]**

| Source | F | df1 | df2 | Sig. | $f^2$ | 95% CI $f^2$ | | size |
|---|---|---|---|---|---|---|---|---|
| Corrected Model | 62.670 | 9 | 189 | 0.000000000 | 2.98 | 2.21 | 3.76 | H |
| SCC | 0.139 | 1 | 92 | 0.709870022 | | | | |
| **GS** | **5.120** | **1** | **84** | **0.026247110** | 0.06 | 0.00 | 0.21 | S |
| **LIQUID** | **97.461** | **2** | **102** | **0.000000000** | 1.91 | 1.22 | 2.72 | H |
| SCC * GS | 0.028 | 1 | 61 | 0.867849722 | | | | |
| SCC * LIQUID | 0.114 | 2 | 175 | 0.892186153 | | | | |
| **GS * LIQUID** | **8.403** | **2** | **179** | **0.000325070** | 0.09 | 0.02 | 0.20 | S |

Probability distribution: Normal Link function: Power(.25)

a. Target: AMT_SUM

**3-WAY ONLY**

**Model Summary**

| | | |
|---|---|---|
| Target | | AMT_SUM |
| Probability Distribution | | Normal |
| Link Function | | Power(.25) |
| Information Criterion | Akaike Corrected | 57.054 |
| | Bayesian | 73.318 |

Information criteria are based on the -2 log likelihood (48.962) and are used to compare models. Models with smaller information criterion values fit better.

**Fixed Effects[a]**

| Source | F | df1 | df2 | Sig. | $f^2$ | 95% CI $f^2$ | | size |
|---|---|---|---|---|---|---|---|---|
| Corrected Model | 57.818 | 11 | 197 | 0.000000000 | 3.23 | 2.42 | 4.03 | H |
| **GS * SCC * LIQUID** | **57.818** | **11** | **197** | **0.000000000** | 3.23 | 2.42 | 4.03 | H |

Probability distribution: Normal Link function: Power(.25)

a. Target: AMT_SUM

**FULL FACTORIAL with 4-LEVEL GENOTYPE (4CoreGT)**

**Model Summary**

| | | |
|---|---|---|
| Target | | AMT_SUM |
| Probability Distribution | | Normal |
| Link Function | | Power(.25) |
| Information Criterion | Akaike Corrected | 57.054 |
| | Bayesian | 73.318 |

Information criteria are based on the -2 log likelihood (48.962) and are used to compare models. Models with smaller information criterion values fit better.

**Fixed Effects[a]**

| Source | F | df1 | df2 | Sig. | $f^2$ | 95% CI $f^2$ | | size |
|---|---|---|---|---|---|---|---|---|
| Corrected Model | 57.818 | 11 | 205 | 0.000000000 | 3.10 | 2.33 | 3.87 | H |
| 4CoreGT | 1.915 | 3 | 81 | 0.133673013 | | | | |
| **LIQUID** | **124.910** | **2** | **131** | **0.000000000** | 1.91 | 1.29 | 2.61 | H |
| **4CoreGT* LIQUID** | **4.373** | **6** | **230** | **0.000332584** | 0.11 | 0.02 | 0.20 | S |

Probability distribution: Normal Link function: Power(.25)

a. Target: AMT_SUM

**MAIN EFFECTS ONLY with 4-LEVEL GENOTYPE (4CoreGT)**

**Model Summary**

| | | |
|---|---|---|
| Target | | AMT_SUM |
| Probability Distribution | | Normal |
| Link Function | | Power(.25) |
| Information Criterion | Akaike Corrected | 44.054 |
| | Bayesian | 60.373 |

Information criteria are based on the -2 log likelihood (35.963) and are used to compare models. Models with smaller information criterion values fit better.

**Fixed Effects[a]**

| Source | F | df1 | df2 | Sig. | $f^2$ | 95% CI $f^2$ | | size |
|---|---|---|---|---|---|---|---|---|
| Corrected Model | 57.988 | 5 | 178 | 0.000000000 | 1.63 | 1.14 | 2.15 | H |
| 4CoreGT | 1.385 | 3 | 130 | 0.250341835 | | | | |
| **LIQUID** | **116.777** | **2** | **197** | **0.000000000** | 1.19 | 0.83 | 1.59 | H |

Probability distribution: Normal Link function: Power(.25)

a. Target: AMT_SUM

### Case Processing Summary

| | N | Percent |
|---|---|---|
| Included | 453 | 100.0% |
| Excluded | 0 | 0.0% |
| Total | 453 | 100.0% |

### Covariance Parameters Summary

| | | |
|---|---|---|
| Covariance Parameters | Residual Effect | 3 |
| | Random Effects | 1 |
| Design Matrix Columns | Fixed Effects | 36 |
| | Random Effects | 1[a] |
| Common Subjects | | 29 |

Common subjects are based on the subject specifications for the residual and random effects and are used to chunk the data for better performance.

a. This is the number of columns per common subject.

### Residual Effect

| Residual Effect | Estimate | Std. Error | Z | Sig. | 95% Confidence Interval Lower | 95% Confidence Interval Upper |
|---|---|---|---|---|---|---|
| Var(LIQUID=1) | 4.792 | 0.591 | 8.103 | **0.000000000** | 3.763 | 6.104 |
| Var(LIQUID=2) | 0.932 | 0.111 | 8.375 | **0.000000000** | 0.737 | 1.177 |
| Var(LIQUID=3) | 0.413 | 0.050 | 8.349 | **0.000000000** | 0.327 | 0.523 |

Covariance Structure: Diagonal    Subject Specification: LITTER * SUBJECT

### Random Effect Block 1

| Random Effect Block | Intercept |
|---|---|
| Intercept | 0.005 |

Covariance Structure: Variance components
Subject Specification: LITTER

### Random Effect

| Random Effect Covariance | Estimate | Std. Error | Z | Sig. | 95% Confidence Interval Lower | 95% Confidence Interval Upper |
|---|---|---|---|---|---|---|
| Var(Intercept) | 0.005 | 0.002 | 2.122 | **0.033845302** | 0.002 | 0.013 |

Covariance Structure: Variance components    Subject Specification: LITTER

### Estimates

| LIQUID | Mean | Std. Error | 95% Confidence Interval Lower | 95% Confidence Interval Upper |
|---|---|---|---|---|
| 1 | 2.872 | 0.234 | 2.434 | 3.367 |
| 2 | 1.074 | 0.085 | 0.913 | 1.255 |
| 3 | 0.653 | 0.052 | 0.553 | 0.766 |

### Estimates

| SCC | LIQ | Mean | Std. Error | 95% Confidence Interval Lower | 95% Confidence Interval Upper |
|---|---|---|---|---|---|
| XX | 1 | 2.685 | 0.319 | 2.105 | 3.378 |
| | 2 | 1.033 | 0.108 | 0.834 | 1.266 |
| | 3 | 0.643 | 0.076 | 0.505 | 0.807 |
| XY | 1 | 3.069 | 0.307 | 2.503 | 3.726 |
| | 2 | 1.116 | 0.124 | 0.890 | 1.382 |
| | 3 | 0.664 | 0.065 | 0.543 | 0.805 |

### Estimates

| GS | LIQ | Mean | Std. Error | 95% Confidence Interval Lower | 95% Confidence Interval Upper |
|---|---|---|---|---|---|
| O | 1 | 2.086 | 0.265 | 1.610 | 2.661 |
| | 2 | 1.016 | 0.117 | 0.803 | 1.270 |
| | 3 | 0.661 | 0.085 | 0.506 | 0.848 |
| T | 1 | 3.862 | 0.342 | 3.220 | 4.596 |
| | 2 | 1.133 | 0.118 | 0.917 | 1.386 |
| | 3 | 0.646 | 0.065 | 0.526 | 0.786 |

### Estimates

| GS | SCC | LIQ | Mean | Std. Error | 95% Confidence Interval Lower | 95% Confidence Interval Upper |
|---|---|---|---|---|---|---|
| O | XX | 1 | 1.618 | 0.337 | 1.047 | 2.396 |
| | | 2 | 1.000 | 0.186 | 0.678 | 1.425 |
| | | 3 | 0.724 | 0.149 | 0.468 | 1.075 |
| | XY | 1 | 2.649 | 0.397 | 1.944 | 3.530 |
| | | 2 | 1.032 | 0.145 | 0.774 | 1.351 |
| | | 3 | 0.601 | 0.092 | 0.439 | 0.804 |
| T | XX | 1 | 4.209 | 0.559 | 3.184 | 5.464 |
| | | 2 | 1.066 | 0.124 | 0.841 | 1.334 |
| | | 3 | 0.568 | 0.090 | 0.409 | 0.769 |
| | XY | 1 | 3.537 | 0.335 | 2.921 | 4.246 |
| | | 2 | 1.204 | 0.203 | 0.847 | 1.662 |
| | | 3 | 0.732 | 0.078 | 0.589 | 0.900 |

### Pairwise Contrasts

| LIQ | Contrast Estimate | Std. Error | t | df | Adj. Sig. | 95% Confidence Interval Lower | Upper | d | 95% CI *d* | | size |
|---|---|---|---|---|---|---|---|---|---|---|---|
| 1 - 2 | 1.799 | 0.203 | 8.847 | 78 | **0.000000000** | 1.335 | 2.262 | 1.00 | 0.73 | 1.27 | H |
| 1 - 3 | 2.219 | 0.219 | 10.138 | 88 | **0.000000000** | 1.686 | 2.752 | 1.08 | 0.82 | 1.34 | H |
| 2 - 3 | 0.420 | 0.072 | 5.848 | 283 | **0.000000014** | 0.279 | 0.562 | 0.35 | 0.23 | 0.47 | S |

The sequential Sidak adjusted significance level is .05. Confidence interval bounds are approximate.

### Simple Contrasts

| LIQ | GS | Contrast Estimate | Std. Error | t | df | Adj. Sig. | 95% Confidence Interval Lower | Upper | d | 95% CI *d* | | size |
|---|---|---|---|---|---|---|---|---|---|---|---|---|
| 1 | O - T | -1.776 | 0.397 | -4.470 | 66 | **0.000031361** | -2.569 | -0.983 | 1.10 | 0.58 | 1.61 | H |
| 2 | O - T | -0.117 | 0.163 | -0.718 | 87 | **0.474583463** | -0.441 | 0.207 | | | | |
| 3 | O - T | 0.014 | 0.110 | 0.130 | 64 | **0.896935227** | -0.206 | 0.235 | | | | |

The sequential Sidak adjusted significance level is .05. Confidence interval bounds are approximate.

### Pairwise Contrasts

| GS | LIQ | Contrast Estimate | Std. Error | t | df | Adj. Sig. | 95% Confidence Interval Lower | Upper | d | 95% CI *d* | | size |
|---|---|---|---|---|---|---|---|---|---|---|---|---|
| O | 1 - 2 | 1.070 | 0.195 | 5.476 | 172 | **0.000000456** | 0.599 | 1.541 | 0.42 | 0.26 | 0.57 | S |
| | 1 - 3 | 1.426 | 0.265 | 5.374 | 141 | **0.000000621** | 0.826 | 2.025 | 0.45 | 0.28 | 0.63 | S |
| | 2 - 3 | 0.356 | 0.106 | 3.344 | 245 | **0.000955315** | 0.146 | 0.565 | 0.21 | 0.09 | 0.34 | S |
| T | 1 - 2 | 2.729 | 0.333 | 8.194 | 53 | **0.000000000** | 1.962 | 3.496 | 1.13 | 0.78 | 1.47 | H |
| | 1 - 3 | 3.216 | 0.308 | 10.428 | 55 | **0.000000000** | 2.457 | 3.976 | 1.41 | 1.03 | 1.77 | H |
| | 2 - 3 | 0.487 | 0.111 | 4.379 | 219 | **0.000018460** | 0.268 | 0.706 | 0.30 | 0.16 | 0.43 | S |

The sequential Sidak adjusted significance level is .05. Confidence interval bounds are approximate.

### Simple Contrasts

| GS | LIQ | SCC | Contrast Estimate | Std. Error | t | df | Adj. Sig. | 95% Confidence Interval Lower | Upper | d | 95% CI *d* | | size |
|---|---|---|---|---|---|---|---|---|---|---|---|---|---|
| O | 1 | XX - XY | -1.031 | 0.516 | -1.997 | 68 | **0.049830240** | -2.060 | -0.001 | 0.48 | 0.00 | 0.96 | S |
| | 2 | XX - XY | -0.032 | 0.238 | -0.135 | 58 | **0.892742688** | -0.508 | 0.443 | | | | |
| | 3 | XX - XY | 0.123 | 0.174 | 0.707 | 47 | **0.483284214** | -0.228 | 0.474 | | | | |
| T | 1 | XX - XY | 0.672 | 0.598 | 1.124 | 46 | **0.266911557** | -0.531 | 1.875 | | | | |
| | 2 | XX - XY | -0.137 | 0.237 | -0.581 | 87 | **0.562825891** | -0.608 | 0.333 | | | | |
| | 3 | XX - XY | -0.164 | 0.112 | -1.471 | 364 | **0.142165565** | -0.384 | 0.055 | | | | |

The sequential Sidak adjusted significance level is .05. Confidence interval bounds are approximate.

### Simple Contrasts

| SCC | LIQ | GS | Contrast Estimate | Std. Error | t | df | Adj. Sig. | 95% Confidence Interval Lower | Upper | d | 95% CI *d* | | size |
|---|---|---|---|---|---|---|---|---|---|---|---|---|---|
| XX | 1 | O - T | -2.591 | 0.647 | -4.002 | 37 | **0.000285746** | -3.902 | -1.280 | 1.32 | 0.60 | 2.02 | H |
| | 2 | O - T | -0.066 | 0.233 | -0.284 | 82 | **0.777435150** | -0.529 | 0.397 | | | | |
| | 3 | O - T | 0.156 | 0.189 | 0.828 | 39 | **0.412519616** | -0.226 | 0.538 | | | | |
| XY | 1 | O - T | -0.888 | 0.423 | -2.099 | 169 | **0.037336054** | -1.724 | -0.053 | 0.32 | 0.02 | 0.63 | S |
| | 2 | O - T | -0.171 | 0.248 | -0.689 | 76 | **0.492983212** | -0.666 | 0.324 | | | | |
| | 3 | O - T | -0.131 | 0.112 | -1.172 | 347 | **0.242003627** | -0.351 | 0.089 | | | | |

The sequential Sidak adjusted significance level is .05. Confidence interval bounds are approximate.

### Pairwise Contrasts

| GS | SCC | LIQ | Contrast Estimate | Std. Error | t | df | Adj. Sig. | 95% Confidence Interval Lower | Upper | d | 95% CI *d* | | size |
|---|---|---|---|---|---|---|---|---|---|---|---|---|---|
| O | XX | 1 - 2 | 0.618 | 0.247 | 2.498 | 237 | **0.035292838** | 0.032 | 1.204 | 0.16 | 0.03 | 0.29 | S |
| | | 1 - 3 | 0.894 | 0.352 | 2.542 | 174 | **0.035292838** | 0.046 | 1.742 | 0.19 | 0.04 | 0.34 | S |
| | | 2 - 3 | 0.276 | 0.165 | 1.677 | 210 | **0.095027487** | -0.048 | 0.600 | 0.12 | -0.02 | 0.25 | S |
| | XY | 1 - 2 | 1.616 | 0.279 | 5.787 | 91 | **0.000000304** | 0.937 | 2.296 | 0.61 | 0.38 | 0.83 | M |
| | | 1 - 3 | 2.048 | 0.369 | 5.546 | 81 | **0.000000720** | 1.206 | 2.889 | 0.62 | 0.38 | 0.85 | M |
| | | 2 - 3 | 0.431 | 0.142 | 3.030 | 230 | **0.002724710** | 0.151 | 0.712 | 0.20 | 0.07 | 0.33 | S |
| T | XX | 1 - 2 | 3.143 | 0.557 | 5.641 | 33 | **0.000005801** | 1.837 | 4.449 | 0.98 | 0.56 | 1.39 | H |
| | | 1 - 3 | 3.641 | 0.521 | 6.991 | 34 | **0.000000141** | 2.333 | 4.949 | 1.20 | 0.75 | 1.63 | H |
| | | 2 - 3 | 0.498 | 0.153 | 3.258 | 350 | **0.001232166** | 0.197 | 0.799 | 0.17 | 0.07 | 0.28 | S |
| | XY | 1 - 2 | 2.334 | 0.306 | 7.636 | 426 | **0.000000000** | 1.648 | 3.020 | 0.37 | 0.27 | 0.47 | S |
| | | 1 - 3 | 2.805 | 0.327 | 8.580 | 242 | **0.000000000** | 2.019 | 3.591 | 0.55 | 0.42 | 0.69 | M |
| | | 2 - 3 | 0.471 | 0.191 | 2.470 | 85 | **0.015518103** | 0.092 | 0.851 | 0.27 | 0.05 | 0.48 | S |

The sequential Sidak adjusted significance level is .05. Confidence interval bounds are approximate.
