## Additional File 4: Supplementary Results Ancillary Stats Example for "Gonadal Sex and Sex-Chromosome Complement Interact to Affect Ethanol Consumption in Adolescent Four Core Genotypes Mice"

**Additional File 4 (Supplementary Results –Ancillary Stats Example.pdf):**
**Includes example SPSS statistical output of bootstrapped ANOVA analysis of mean pre-session body weight.**

| KEY | | |
|---|---|---|
| SCC | = | Sex Chromosome Complement |
| GS | = | Gonadal Sex |
| LIQ | = | Liquid |
| O | = | Ovaries |
| T | = | Testes |
| 1 (LIQ) | = | 20% EtOH |
| 2 (LIQ) | = | 10% EtOH |
| 3 (LIQ) | = | 0% EtOH |
| BW PRE | = | Pre-session body weight (g) |
| BECs | = | Blood Ethanol Concentrations (md/dL) |
| bANOVA | = | bootstrapped ANalysis Of VAriance |
| PH | = | post hoc paired-means comparisons |
| d | = | Cohen's *d* |
| $f^2$ | = | Cohen's *f* |

### FULL FACTORIAL

#### Tests of Between-Subjects Effects

Dependent Variable: BW PRE (g; xS)

| Source | | Type III Sum of Squares | df | Mean Square | F | Sig. | Partial Eta Squared | $f^2$ | 95% CI $f^2$ | | size |
|---|---|---|---|---|---|---|---|---|---|---|---|
| Intercept | Hypothesis | 27263.287 | 1 | 27263.287 | 6728.679 | 0.000000000 | 0.995 | 215.84 | 121.94 | 336.04 | H |
| | Error | 126.312 | 31.174 | 4.052[a] | | | | | | | |
| LITTER | Hypothesis | 136.253 | 28 | 4.866 | 4.819 | 0.000000001 | 0.531 | 1.13 | 0.49 | 1.42 | H |
| | Error | 120.154 | 119 | 1.010[b] | | | | | | | |
| SCC | Hypothesis | 12.342 | 1 | 12.342 | 12.223 | 0.000664088 | 0.093 | 0.10 | 0.02 | 0.25 | S |
| | Error | 120.154 | 119 | 1.010[b] | | | | | | | |
| GS | Hypothesis | 77.503 | 1 | 77.503 | 76.759 | 0.000000000 | 0.392 | 0.65 | 0.35 | 1.02 | H |
| | Error | 120.154 | 119 | 1.010[b] | | | | | | | |
| SCC * GS | Hypothesis | 0.030 | 1 | 0.030 | 0.030 | 0.863643781 | 0.000 | | | | |
| | Error | 120.154 | 119 | 1.010[b] | | | | | | | |

a. .789 MS(LITTER) + .211 MS(Error)

b. MS(Error)

### SIMPLE EFFECT: SCC

#### Univariate Tests

Dependent Variable: BW PRE (g; xS)

| GS | SCC | Sum of Squares | df | Mean Square | F | Sig. | Partial Eta Squared | $f^2$ | 95% CI $f^2$ | | size |
|---|---|---|---|---|---|---|---|---|---|---|---|
| F | Contrast | 6.971 | 1 | 6.971 | 6.904 | 0.009730649 | 0.055 | 0.06 | 0.00 | 0.18 | S |
| | Error | 120.154 | 119 | 1.010 | | | | | | | |
| M | Contrast | 5.929 | 1 | 5.929 | 5.872 | 0.016889634 | 0.047 | 0.05 | 0.00 | 0.16 | S |
| | Error | 120.154 | 119 | 1.010 | | | | | | | |

Each F tests the simple effects of SCC within each level combination of the other effects shown. These tests are based on the linearly independent pairwise comparisons among the estimated marginal means.

### SIMPLE EFFECT: GS

#### Univariate Tests

Dependent Variable: BW PRE (g; xS)

| SCC | GS | Sum of Squares | df | Mean Square | F | Sig. | Partial Eta Squared | $f^2$ | 95% CI $f^2$ | | size |
|---|---|---|---|---|---|---|---|---|---|---|---|
| XX | Contrast | 40.195 | 1 | 40.195 | 39.809 | 0.000000005 | 0.251 | 0.34 | 0.15 | 0.60 | M |
| | Error | 120.154 | 119 | 1.010 | | | | | | | |
| XY | Contrast | 41.611 | 1 | 41.611 | 41.212 | 0.000000003 | 0.257 | 0.35 | 0.16 | 0.62 | H |
| | Error | 120.154 | 119 | 1.010 | | | | | | | |

Each F tests the simple effects of GS within each level combination of the other effects shown. These tests are based on the linearly independent pairwise comparisons among the estimated marginal means.

### MAIN EFFECTS ONLY

#### Tests of Between-Subjects Effects

Dependent Variable: BW PRE (g; xS)

| Source | | Type III Sum of Squares | df | Mean Square | F | Sig. | Partial Eta Squared | $f^2$ | 95% CI $f^2$ | | size |
|---|---|---|---|---|---|---|---|---|---|---|---|
| Intercept | Hypothesis | 27264.918 | 1 | 27264.918 | 6757.822 | 0.000000000 | 0.995 | 216.39 | 122.32 | 336.77 | H |
| | Error | 125.998 | 31.230 | 4.035[a] | | | | | | | |
| LITTER | Hypothesis | 136.283 | 28 | 4.867 | 4.860 | 0.000000001 | 0.531 | 1.13 | 0.50 | 1.42 | H |
| | Error | 120.184 | 120 | 1.002[b] | | | | | | | |
| SCC | Hypothesis | 12.328 | 1 | 12.328 | 12.309 | 0.000635312 | 0.093 | 0.10 | 0.02 | 0.25 | S |
| | Error | 120.184 | 120 | 1.002[b] | | | | | | | |
| GS | Hypothesis | 77.474 | 1 | 77.474 | 77.355 | 0.000000000 | 0.392 | 0.64 | 0.36 | 1.01 | H |
| | Error | 120.184 | 120 | 1.002[b] | | | | | | | |

a. .785 MS(LITTER) + .215 MS(Error)

b. MS(Error)

### INTERACTION ONLY

#### Tests of Between-Subjects Effects

Dependent Variable: BW PRE (g; xS)

| Source | | Type III Sum of Squares | df | Mean Square | F | Sig. | Partial Eta Squared | $f^2$ | 95% CI $f^2$ | | size |
|---|---|---|---|---|---|---|---|---|---|---|---|
| Intercept | Hypothesis | 27263.287 | 1 | 27263.287 | 6728.679 | 0.000000000 | 0.995 | 215.84 | 121.94 | 336.04 | H |
| | Error | 126.312 | 31.174 | 4.052[a] | | | | | | | |
| LITTER | Hypothesis | 136.253 | 28 | 4.866 | 4.819 | 0.000000001 | 0.531 | 1.13 | 0.49 | 1.42 | H |
| | Error | 120.154 | 119 | 1.010[b] | | | | | | | |
| SCC * GS | Hypothesis | 87.283 | 3 | 29.094 | 28.815 | 0.000000000 | 0.421 | 0.73 | 0.40 | 1.11 | H |
| | Error | 120.154 | 119 | 1.010[b] | | | | | | | |

a. .789 MS(LITTER) + .211 MS(Error)

b. MS(Error)

**Between-Subjects Factors**

| | | N |
|---|---|---|
| SCC | XX | 78 |
| | XY | 73 |
| GS | O | 77 |
| | T | 74 |
| LITTER | 107 | 3 |
| | 114 | 2 |
| | 117 | 6 |
| | 119 | 6 |
| | 122 | 7 |
| | 128 | 6 |
| | 131 | 4 |
| | 134 | 4 |
| | 136 | 6 |
| | 144 | 5 |
| | 152 | 7 |
| | 156 | 7 |
| | 158 | 8 |
| | 168 | 10 |
| | 169 | 8 |
| | 175 | 4 |
| | 180 | 7 |
| | 186 | 8 |
| | 188 | 5 |
| | 189 | 5 |
| | 190 | 9 |
| | 51 | 3 |
| | 62 | 4 |
| | 72 | 2 |
| | 76 | 4 |
| | 77 | 1 |
| | 80 | 3 |
| | 84 | 4 |
| | 98 | 3 |

**Bootstrap Specifications**

| | |
|---|---|
| Sampling Method | Simple |
| Number of Samples | 1000 |
| Confidence Interval Level | 95.0% |
| Confidence Interval Type | Percentile |

**Expected Mean Squares[a,b]**

| Source | Variance Component | | |
|---|---|---|---|
| | Var(LITTER) | Var(Error) | Quadratic Term |
| Intercept | 3.976 | 1.000 | Intercept, SCC, GS, SCC * GS |
| LITTER | 5.040 | 1.000 | |
| SCC | 0.000 | 1.000 | SCC, SCC * GS |
| GS | 0.000 | 1.000 | GS, SCC * GS |
| SCC * GS | 0.000 | 1.000 | SCC * GS |
| Error | 0.000 | 1.000 | |

a. For each source, the expected mean square equals the sum of the coefficients in the cells times the variance components, plus a quadratic term involving effects in the Quadratic Term cell.

b. Expected Mean Squares are based on the Type III Sums of Squares.

**Estimates**

Dependent Variable: BW PRE (g; xS)

| SCC | GS | Mean | Std. Error | 95% Confidence Interval | | Bootstrap for Mean[a] | | | |
|---|---|---|---|---|---|---|---|---|---|
| | | | | Lower Bound | Upper Bound | Bias | Std. Error | 95% Confidence Interval | |
| | | | | | | | | Lower | Upper |
| XX | O | 14.897 | 0.181 | 14.539 | 15.254 | -0.030 | 0.175 | 14.513 | 15.221 |
| | T | 16.503 | 0.179 | 16.147 | 16.858 | -0.072 | 0.220 | 16.019 | 16.841 |
| XY | O | 14.221 | 0.191 | 13.842 | 14.600 | -0.018 | 0.177 | 13.866 | 14.559 |
| | T | 15.887 | 0.184 | 15.524 | 16.251 | -0.074 | 0.210 | 15.390 | 16.211 |

a. Unless otherwise noted, bootstrap results are based on 1000 bootstrap samples

### Pairwise Comparisons

Dependent Variable: BW PRE (g; xS)

| GS | SCC | Mean Difference (I-J) | Std. Error | Sig.[b] | 95% Confidence Interval for Difference[b] Lower Bound | Upper Bound |
|---|---|---|---|---|---|---|
| O | XX - XY | .676* | 0.257 | **0.009730649** | 0.167 | 1.185 |
| T | XX - XY | .615* | 0.254 | **0.016889634** | 0.112 | 1.118 |

Based on estimated marginal means

*. The mean difference is significant at the .05 level.

b. Adjustment for multiple comparisons: Sidak.

### Bootstrap for Pairwise Comparisons

Dependent Variable: BW PRE (g; xS)

| GS | SCC | Mean Difference (I-J) | Bootstrap[a] Bias | Std. Error | Sig. (2-tailed) | 95% Confidence Interval Lower | Upper |
|---|---|---|---|---|---|---|---|
| O | XX - XY | 0.676 | -0.012 | 0.223 | **0.005994006** | 0.251 | 1.089 |
| T | XX - XY | 0.615 | 0.001 | 0.315 | **0.056943057** | -0.010 | 1.238 |

a. Unless otherwise noted, bootstrap results are based on 1000 bootstrap samples

### Pairwise Comparisons

Dependent Variable: BW PRE (g; xS)

| SCC | GS | Mean Difference (I-J) | Std. Error | Sig.[b] | 95% Confidence Interval for Difference[b] Lower Bound | Upper Bound |
|---|---|---|---|---|---|---|
| XX | O - T | -1.606* | 0.255 | **0.000000005** | -2.110 | -1.102 |
| XY | O - T | -1.667* | 0.260 | **0.000000003** | -2.181 | -1.153 |

Based on estimated marginal means

*. The mean difference is significant at the .05 level.

b. Adjustment for multiple comparisons: Sidak.

### Bootstrap for Pairwise Comparisons

Dependent Variable: BW PRE (g; xS)

| SCC | GS | Mean Difference (I-J) | Bootstrap[a] Bias | Std. Error | Sig. (2-tailed) | 95% Confidence Interval Lower | Upper |
|---|---|---|---|---|---|---|---|
| XX | O - T | -1.606 | 0.043 | 0.275 | **0.000999001** | -2.086 | -1.021 |
| XY | O - T | -1.667 | 0.056 | 0.264 | **0.000999001** | -2.117 | -1.080 |

a. Unless otherwise noted, bootstrap results are based on 1000 bootstrap samples
